# Combinatorial optimization of protein systems in synthetic cells

**DOI:** 10.64898/2026.02.25.707944

**Authors:** Marijn van den Brink, Nico J. Claassens, Christophe Danelon

## Abstract

*In vitro* reconstitution of protein systems – e.g., metabolic pathways, genetic circuits or biosensors – often requires optimization to enhance their activity. Combinatorial DNA libraries that simultaneously target multiple genes allow for a holistic optimization strategy by studying the interplay between the systems’ components, which may reveal DNA variants that would be hidden when testing each element in isolation. Here, we screen large populations of synthetic vesicles that express combinatorial DNA variants of a DNA self-replicator or a phospholipid synthesis pathway. We simultaneously vary the strengths of multiple RBSs or synonymously mutate the first codons of multiple genes to explore the effects of the protein translation rates directly on the functionality of the two core synthetic cell modules. We isolated high performers through DNA self-selection or functional screening by fluorescence-activated cell sorting. Long-read sequencing of the fittest variants informed on the optimal RBS strengths and base substitutions in the first codons and indicated which genes were most impactful in regulating the functionality of the protein systems. Single-mutation data were used to predict the fitness of combinatorial variants, which was compared with the experimental fitness observed. The theoretical fitness of combinatorial variants was extremely predictive for the two-gene library of the DNA replicator but less for the larger pathway library. Altogether, our approach exemplifies how combinatorial testing can be expanded from single proteins to multiprotein systems, which can in the future be extended to the evolutionary engineering of even larger genetic and metabolic networks, and eventually an entire artificial cell.

**GRAPHICAL ABSTRACT:** 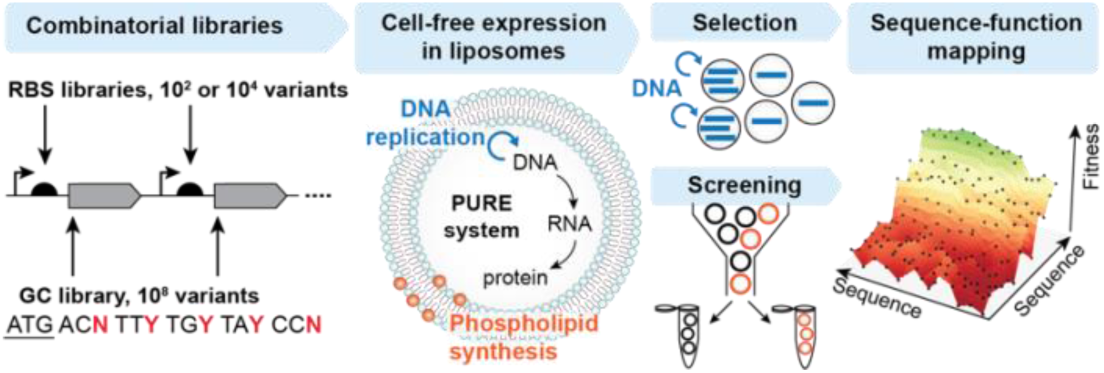

## INTRODUCTION

Cell-free gene expression offers an ideal playground to explore biological designs. It finds applications in prototyping, biomanufacturing, and has emerged as a platform to build a synthetic cell^1,2^. The reconstitution of metabolic pathways, genetic circuits or biosensors often requires optimization steps to improve functional performance. For example, the engineering of regulatory parts, such as promoters, terminators and ribosome binding sites (RBSs) has become a routine method to control gene expression – and thereby protein concentrations – *in vivo* and *in vitro*, thus enhancing a certain objective function, for instance maximizing the yield of a desired enzymatic product^1,3^.

Large libraries of DNA variants can be assayed by high-throughput screening of femto-to picoliter cell-free expression reactions encapsulated in lipid vesicles or droplets^4–13^. While many examples of such *in vitro* evolution screens have been reported for the optimization of single genes, multivariate optimization involving multiple proteins is more challenging due to the vast combinatorial space and unpredictable epistatic effects. Examples of multi-protein system optimization targets include (i) the production of proteins at stoichiometric amounts within a macromolecular complex, (ii) the balanced production of enzymes involved in a pathway to enhance metabolic fluxes and avoid accumulation of (toxic) intermediate products, (iii) fine tuning gene expression profiles for efficient allocation of resources, and (iv) the regulation of genetic networks to promote the integration of and coordinate different biological functions. Living organisms navigate such complexity through natural evolution across extremely long timespans. Accelerating the optimization of multigene functions through directed evolution would require approaches that reduce the problem of combinatorial explosion, for instance by generating ‘small but smart’ libraries, and implement selection procedures that retain the best performing designs^14–16^.

In this study, we performed *in vitro* evolution of two genetic modules, each encoding multiple proteins involved in DNA replication or phospholipid synthesis. The genomes were expressed with PURE system (Protein synthesis Using Recombinant Elements)^17^ in liposome-based synthetic cells. We here define ‘synthetic cell’ as an umbrella term for the compartmentalized expression of genes encoding a cellular function. We investigated the effects of modified RBS sequences or the GC content of the first codons on the functionality of the two genetic systems. It is known from previous studies that translation initiation is often the rate-limiting step in bacterial translation^3,18^, and that the initiation rate also controls protein yield in PURE system^19,20^. However, it remains unclear how changing the translation rates of multiple genes impacts the overall expression pattern in PURE system, possibly distorted by epistatic interactions between genes competing for translation resources (e.g., tRNAs, tRNA synthetases, amino acids). It is also unknown whether control over protein amounts is effective in regulating the functionality of the protein systems in synthetic cells.

To address these questions, we expressed semi-rational, combinatorial DNA libraries in synthetic cells. The fittest variants were isolated either through selection by DNA self-replication or through a FACS-based liposome screening method with a reporter to detect the output molecule of the phospholipid synthesis pathway. Long-read sequencing of the fittest variants informed on the optimal RBS strengths and base substitutions in the first codons. The data also suggested which genes were most impactful in regulating the functionality of the modules, and whether the effects of multiple mutations were predictable or unexpected. Altogether, these results lay the groundwork for future evolutionary engineering efforts by demonstrating how increased DNA template complexity can be harnessed in synthetic cell systems.

## RESULTS

### Design of a combinatorial library of RBS variants in a DNA replicator

Protein-primed DNA replication was established in liposomes by the *in vitro* expression of two proteins from the *Bacillus subtilis* bacteriophage Phi29^21^ (**Fig. 1a**). The Phi29 DNA polymerase (DNAP) and terminal protein (TP) were produced from a linear template flanked with the origins of replication from the Phi29 genome. DNAP and TP form a 1:1 heterodimer and recognize the terminal origins of replication, where the TP forms a covalent bond with the first deoxyadenosine monophosphate (dAMP) of the new 5’ DNA strand^22^. The polymerase then elongates the TP-primed DNA strand with strand-displacement of the original DNA. The process is further facilitated by the addition of purified double-stranded binding protein (DSB) and single-stranded binding protein (SSB) to unwind the DNA at the origins and stabilize displaced single-stranded DNA, respectively.

**Fig. 1.**
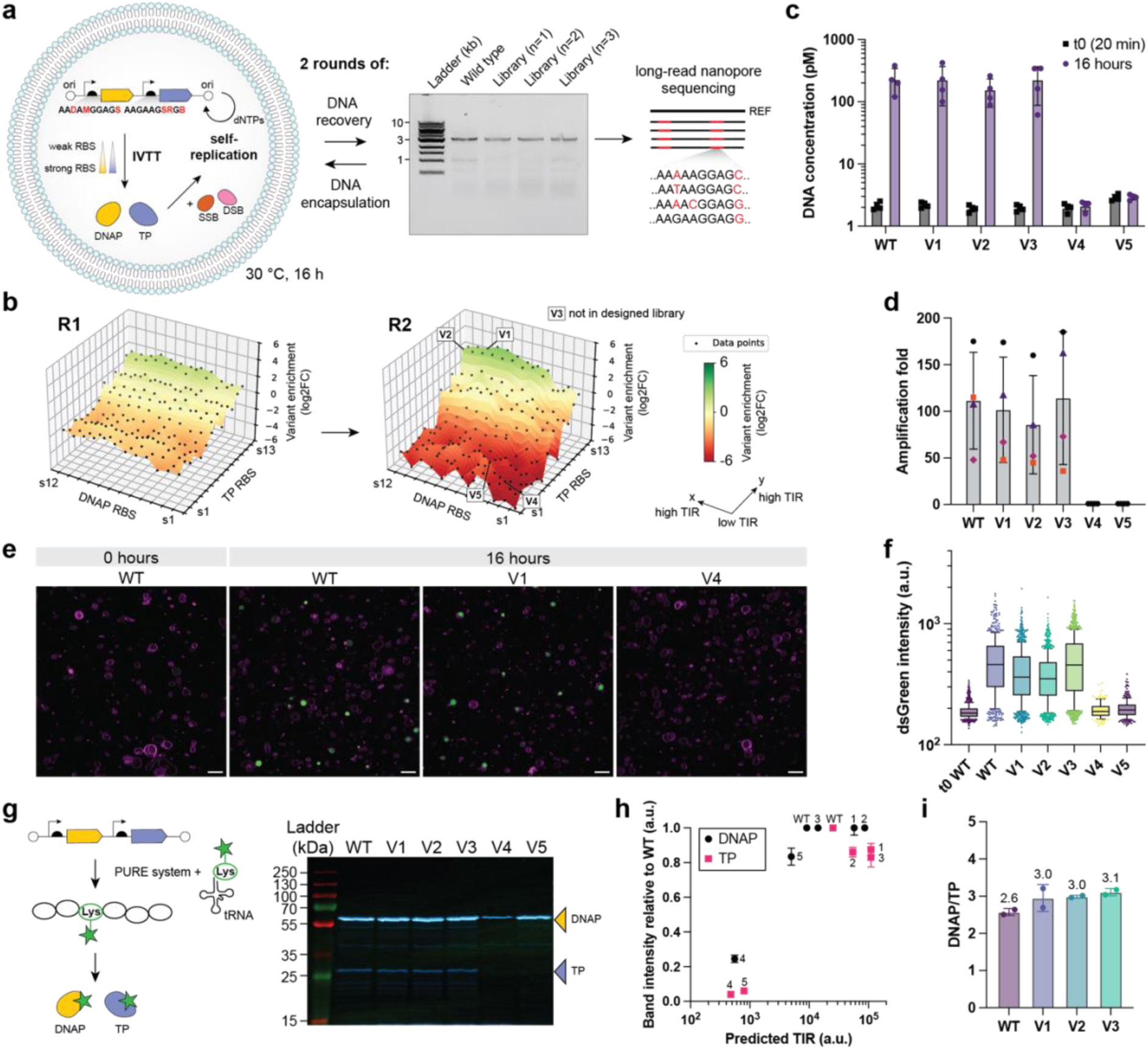
Selection of RBS variants through DNA self-replication. **a** Schematic illustration of *in vitro* transcription-translation (IVTT) and DNA self-replication of the RBS library. DNAP and TP were expressed in liposomes by PURE system. SSB and DSB were supplied as purified proteins. DNA from the liposome samples was recovered by PCR, verified by gel electrophoresis and analyzed by nanopore sequencing. **b** 3D landscapes displaying the log2FC in read frequency of combinatorial variants. The RBS variants on the x- and y-axes are ordered from low-to-high predicted TIR. The landscapes show the log2-transformed average FC of three biological replicates. The number of reads covering the 156 variants are 28,978 in R0 (pre-selection), 10,140 to 16,998 per replicate in R1, and 3,900 to 6,060 per replicate in R2. Data from the individual repeats are shown in Fig. S3. **c** Quantification of DNA concentration by qPCR for clonal DNA variants V1-5 before replication (t0, incubation 20 min) and after replication (16 h) at 30 °C (n=4 biological replicates). **d** Amplification fold of the data in panel c, calculated as the DNA concentration at 16 h divided by the concentration at 20 min. Symbols represent the different biological replicates (n=4). **e** Confocal fluorescence images of liposomes with different clonal DNA variants. Magenta, Cy5-conjugated lipids; green, dsGreen; Scale bars: 10 µm. **f** Quantification of the dsGreen intensity of dsGreen-positive liposomes (>129 a.u.) from confocal microscopy images. Number of dsGreen-positive liposomes: 599 (t0 WT), 487 (WT), 1,056 (V1), 1,040 (V2), 1,255 (V3), 170 (V4), 332 (V5). **g** SDS-PAGE of bulk (no liposomes) PURE reaction samples from clonal DNA variants with GreenLys protein labeling reagent. The fainter bands for TP compared to DNAP could be the result of the lower number of lysine residues of TP (27) compared to DNAP (64). **h** Band intensities from the SDS-PAGE gel in panel g relative to the wild type versus the corresponding predicted TIRs (n=2). The DNA variant is indicated for each data point. **i** Ratio of corrected band intensities for DNAP over TP. Corrected band intensities were calculated by dividing the adjusted band intensity (intensity with background subtracted) by the number of lysines per protein (64 for DNAP, 27 for TP).

The effects of changing the absolute and stoichiometric amounts of DNAP, TP and DNA on replication efficacy has been studied using purified proteins in simple buffer conditions^23–25^. However, it is unknown how the production rates of DNAP and TP impact DNA replication in this reconstituted autocatalytic network, where protein concentrations vary over time and DNA is also the substrate of RNA polymerases which may collide with DNAP^26^. To test a range of protein production levels, we mutated the RBSs for each of the two coding sequences on the template. We targeted the RBSs, because (i) protein production level is better correlated with translation rate than transcription rate in PURE system^19,20^, and (ii) translation initiation is often a major rate-limiting step in protein translation in *Escherichia coli*^18,27^ thus possibly also in PURE system^28^. Because the strength of an RBS is influenced by the ∼35 nucleotides upstream and downstream the start codon^18,29–31^, we designed a specific set of RBS sequences for each of the protein coding sequences using RBS Calculator^18^. The library was designed to maximize the range of predicted translation initiation rates (TIRs) per gene, while keeping the number of mutations between one and five base pair substitutions relative to each wild-type RBS (**Table S1**). The combinatorial library consisted of 12 RBS variants for DNAP and 13 RBS variants for TP, thus in total 156 combinatorial variants. We constructed the library by our recently published protocol for multiplex one-step single-stranded DNA annealing protein (SSAP)-mediated plasmid diversification (MOSAIC)^32^. In short, this method relies on plasmid recombineering in an *E. coli* strain equipped with recombination machinery and requires just one co-electroporation of the target plasmid and degenerate ssDNA oligos encoding the mutations of interest. After one round of MOSAIC, 33% of the DNA library variants harbored two mutated RBSs (**Fig. S1**). The remaining fraction of the library consisted of variants with only one modified RBS (46%) and the wild type (22%).

### Self-selection of RBS variants through DNA replication

To establish a genotype-phenotype link, the library of DNA replicators was expressed by PURE system in liposomes at an average DNA copy number of one per liposome (λ = 1, Methods section). After 16 h of incubation at 30 °C, quantitative PCR (qPCR) analysis confirmed 10 to 100 fold DNA amplification (**Fig. S2a-b**). The resulting population of full-length DNA was amplified from the liposome sample by PCR and re-encapsulated in liposomes for a second round of *in vitro* transcription-translation-replication (IVTTR) (**Fig. 1a**, **Fig. S2c**). Nanopore sequencing of the initial library and the recovered DNA after round one and two revealed changes in combinatorial variant frequencies (**Fig. 1b**). Certain combinations of RBS variants were enriched (log2 fold change (log2FC) > 0), while others decreased in frequency (log2FC < 0) or disappeared from the library (**Fig. 1b**, **Fig. S3**). Overall, the sequence landscape showed a gradual increase of fitness, defined as log2FC, with increasing predicted TIR. The most enriched variants harbored strong RBSs for both TP and DNAP. Interestingly, the fitness was mostly dependent on the predicted TIR of TP and less of DNAP (**Fig. S4**). This could mean that either TP is required in higher amounts and/or the TP RBS library covers a lower expression range than predicted. Specifically, for both DNAP and TP, the log2FC of the RBS variants increased with the predicted TIR up to a certain TIR, whereafter the log2FC plateaued (TP) or slightly decreased (DNAP) (**Fig. S4**). Some RBS variants led to unpredictable results, indicated by a rugged pattern, most pronounced for TP RBSs s8-s11, possibly due to inaccurate predictions of the TIRs for some RBSs (**Fig. S3, S4**). Three biological replicates resulted in similar frequency changes, indicating high reproducibility of the self-selection system behavior (**Fig. S3**). Furthermore, while mutations could be introduced during in-liposome replication by the Phi29 DNA polymerase or during DNA recovery by PCR, no mutations outside the targeted RBS regions were detected in the DNA population with frequency ≥1%, demonstrating that the observed variations in DNA replicator fitness exclusively come from the mutations in the RBS sequences.

### Reverse engineering of clonal self-replicator variants reveals high performance

We then sought to validate that the observed changes in variant frequency were proportional to the ability to self-replicate (i.e., the fitness of each DNA variant). Therefore, we constructed a few clonal variants from the sequence landscape, and we assessed their self-replication ability in liposomes using qPCR (**Table S2, Fig. 1c-d**). Two high-frequency variants, V1 and V2, showed high replication folds. In contrast, V4 and V5, two variants in the “valley” of the sequence landscape, did not replicate at all. We tested another DNA variant, V3, which had the same TP RBS as V1 but a DNAP RBS that was not part of the theoretical library. This DNAP RBS variant was most likely generated as by-product during library generation with MOSAIC or linearization of the library with PCR, as it was already present before encapsulation in liposomes. V3 started at a 50-fold lower frequency than V1 in the initial library (R0), but V3 was enriched 3.2 ± 0.5 fold more than V1 in R2 (n=3). Indeed, clonal V3 also replicated well (**Fig. 1c-d**).

In addition to the population-level qPCR analysis, we then investigated the performance of each replicator variant at the single-liposome level using confocal fluorescence microscopy (**Fig. 1e**). We detected DNA-replicating liposomes based on the intensity of the fluorescent dye dsGreen, which binds specifically to double-stranded DNA. We plotted the average dsGreen intensity per liposome (**Fig. 1f, Fig. S5**) and found similar results as the qPCR data, corroborating that the variant frequency change (log2FC) is indeed an accurate measure for its fitness.

Lastly, we directly measured the amounts of synthesized proteins from the different RBS variants to examine the correlation with fitness. The proteins were expressed in bulk reactions in the presence of tRNA preloaded with fluorescently labelled lysine, and we quantified the fluorescence intensity of the bands after SDS-PAGE (**Fig. 1g-h**, **Fig. S6**). V1-3 showed clear bands for DNAP and TP, with intensities similar to the wild type, despite having higher predicted TIR values for both DNAP (14,117-86,579 a.u.) and TP (54,294-110,313) (comparison to wild type (WT): DNAP: 9,123; TP: 25,228). This observation coincides with the plateauing fitness (log2FC) at high predicted TIRs for DNAP and TP in the library dataset (**Fig. S4**), and with the similar DNA amplification folds for the high-TIR clonal variants (**Fig. 1d-f**). The estimated ratio of molecules DNAP over TP, derived by normalizing the band intensity to the number of lysine residues per protein, increased slightly from 2.6 for the wild type to 3.0-3.1 for the library variants V1-3 (**Fig. 1i**). In contrast, V4 and V5, which have lower TIRs for both DNAP and TP, showed reduced DNAP and TP expression levels, most likely impairing their inability to replicate. Notably, we here quantified the end-point expression level, which may differ from the protein yield and stoichiometry at earlier time points in the reaction.

### Design of a combinatorial RBS library of a 4-gene phospholipid synthesis pathway

Next, we performed multivariate optimization of a four-gene phospholipid synthesis pathway, generating a cascade of reactions catalyzed by membrane-associated or transmembrane proteins (**Fig. 2a**). We selected the four upstream enzymes of the *E. coli* Kennedy phospholipid pathway PlsB, PlsC, CdsA and PssA. These enzymes stepwise convert oleoyl-coenzyme A (oleoyl-CoA) and glycerol-3-phosphate (G3P) to phosphatidylserine (PS) in the presence of the co-substrates cytidine triphosphate (CTP) and L-serine. The amount of final product PS is positively correlated with the fluorescence intensity of the PS-specific reporter LactC2-mCherry, enabling the quantitative assessment of PS production and membrane incorporation at the single-liposome level^33^.

**Fig. 2.**
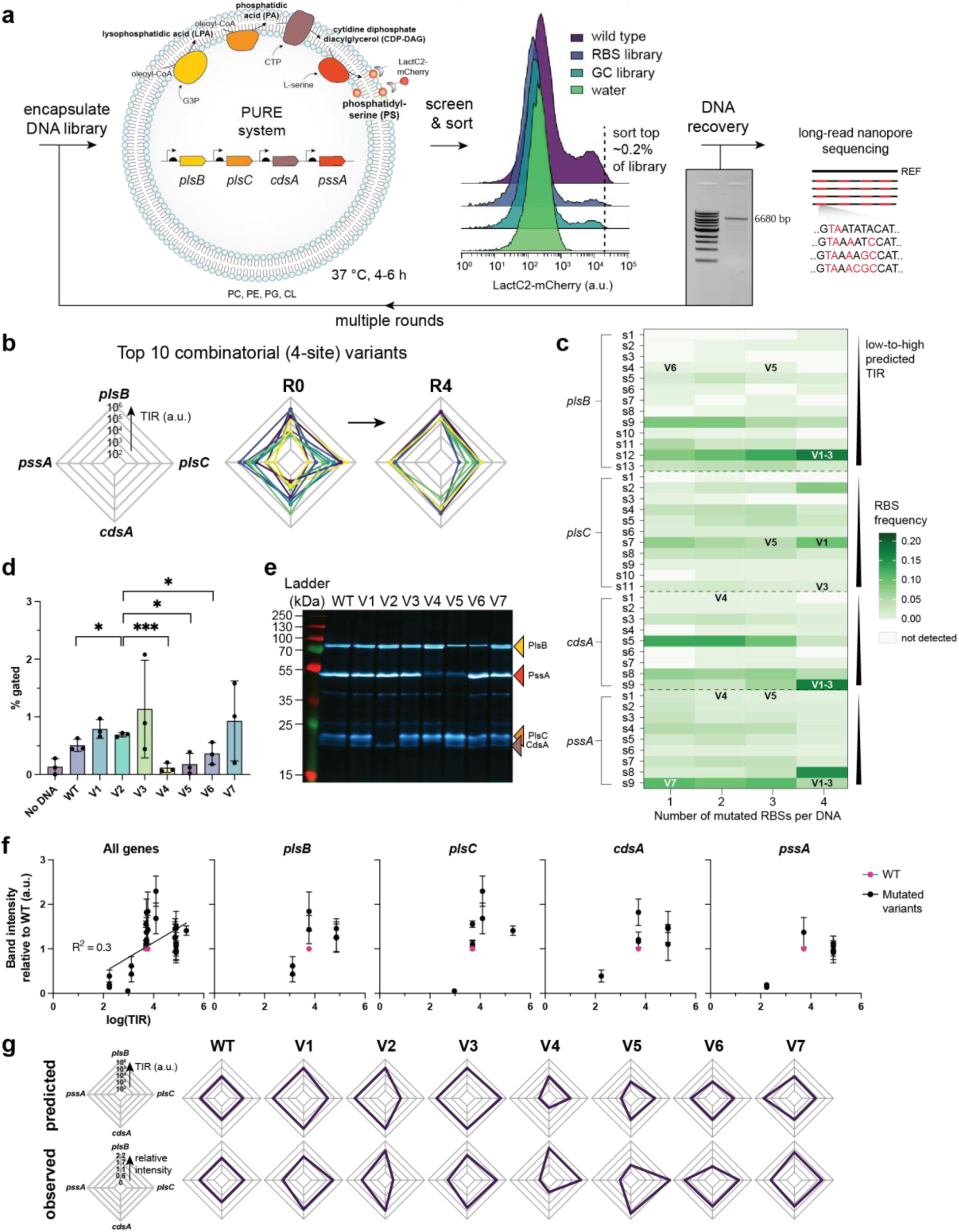
Activity-based selection of DNA designs for PS production from a library of four-site combinatorial variants. **a** Schematic illustration of the workflow. Liposomes express four enzymes from a DNA template with diversified RBSs. The enzymes form a pathway to produce PS, measured by the LactC2-mCherry probe. Four subsequent rounds of sorting the liposomes with highest LactC2-mCherry signal and re-encapsulation of the recovered DNA were performed prior to long-read sequencing. **b** Radar charts displaying the top 10 combinatorial variants with the highest frequency in the initial library (R0) and after four rounds of sorting. The variants are listed in Table S6. **c** Heatmap displaying the frequency of each RBS variant. The frequency was calculated relative to the library fractions with *N* number of mutated RBSs per DNA. Number of reads per library fraction (each column in the heatmap): 9,123 (1-site), 8,510 (2-site), 3,119 (3-site), 697 (4-site). **d** Analysis of clonal DNA variants from the RBS library by flow cytometry. The percentage of gated liposomes was calculated using a similar gate as the one used for sorting the DNA library. Unpaired *t*-test: * *P* ≤ 0.05; *** *P* ≤ 0.001. n = 3 biological replicates. **e** SDS-PAGE with bulk PURE system reactions (no liposomes) containing the GreenLys reagent for co-translational labeling of proteins expressed from clonal DNA variants. **f** Band intensities were measured from the SDS-PAGE gel in panel g, normalized to the wild type, and plotted against the corresponding predicted TIRs (n=2). The appended curve is the linear regression for all genes combined with R^2^ = 0.3, *P* < 0.0001. **g** Radar charts displaying the predicted TIRs and observed relative protein expression levels.

While *in vivo*, the phospholipid synthesis pathway is mainly regulated post-translationally by an (unknown) allosteric regulator of PlsB activity^34^ and metabolic regulation of phosphatidylglycerol (PG) and phosphatidylethanolamine (PE)^35^, we reasoned that a minimal synthetic approach based on translational control could modulate phospholipid production *in vitro*. We hypothesized that the yield of PS production depends on the stoichiometry of the expression of the four enzymes, as observed in other enzymatic cascades^14,36,37^. Previously, it was found that phosphatidic acid (PA), the product of PlsC, significantly accumulates in the Kennedy pathway reconstituted in liposomes, suggesting potential for further optimization^33^. However, the yield of PS has a nontrivial dependency on the four RBS sequences, as the rate-limiting step is unknown and unexpected effects of mutating RBSs could arise from protein folding, membrane insertion/binding, allosteric regulation, inhibitory effects, or resource competition.

To tackle this multivariate problem, we generated a 4-site combinatorial RBS library by MOSAIC consisting of 13 (PlsB) × 11 (PlsC) × 9 (CdsA) × 9 (PssA) = 11,583 variants (**Table S3**). With a MOSAIC editing efficiency of ∼50% per target locus, we obtained a library that consisted for 11% of variants with all RBSs mutated, and the remainder of the library consisted of library fractions where zero, one, two or three RBSs were mutated (**Fig. S7a, c**). Using nanopore sequencing, we detected all designed variants that had one or two mutated RBSs, and 84% and 35% of all designed variants with three and four mutated RBSs, respectively, but more of the designed variants are likely covered in the generated library as suggested by rarefaction plots (**Fig. S8**). Hereafter, we name the fractions of the library with *N* mutated target loci (RBSs) “*N*-site fraction” (e.g., 1-site fraction).

### Enrichment of library variants with high PS yield through screening by FACS

We encapsulated the linearized library in liposomes at an average DNA copy number per liposome λ = 0.2 (Methods). Functionality screening of the liposomes was performed by measuring the fluorescence intensity of the PS probe LactC2-mCherry with flow cytometry. We typically screened ∼3-6 million liposomes of which 2-13% showed PS production (**Fig. S9-S10**). We calculated the screening coverage of the DNA library as shown in **Table S4**, taking into account that 13 ± 8% of a clonal liposome population is active (i.e., the performance of the wild type, **Fig. S9a**, n=5 biological replicates). The screening coverage was approximately 3×10^3^-fold for the 1-site fraction (42 variants), 3×10^2^-fold for the 2-site fraction (656 variants), 3×10^1^-fold for the 3-site fraction (4,518 variants) and 6-fold of the 4-site combinatorial library fraction (11,583 DNA variants). Using FACS, we sorted the liposomes with the highest LactC2-mCherry intensity, less than 1% of all liposomes, corresponding to 10,000-30,000 liposomes. We amplified the DNA from the sorted liposome sample by PCR and analyzed the DNA sequences by nanopore sequencing. We performed four subsequent rounds of sorting, DNA amplification and re-encapsulation in liposomes.

The combinatorial (4-site) genotypes of the top ten most frequent DNA variants are presented in **Fig. 2b** (and **Fig. S12**). Due to the relatively low sequencing coverage of the 3-site and 4-site library fractions in the presort library (R0) (on average 4 ± 4 (3-site) and 2 ± 2 (4-site) reads per variant), we used the read frequency as a measure of fitness instead of calculating the fold change in frequency relative to R0. Comparing the initial library to that after four rounds of sorting, we see a strong shift towards high protein expression levels for PlsB, CdsA and PssA, while a range of RBS strengths were present for PlsC. To obtain a more holistic view on the full dataset, we then plotted the average frequency of each RBS for each fraction of the library (**Fig. 2c, S12**). These results support the observation that high-fitness variants contain *plsB*, *cdsA* and *pssA* RBSs with high predicted TIRs (*plsB*: 21,409-112,986 a.u.; *cdsA*/*pssA*: 31,879-78,824 a.u.) but a more even distribution of frequencies is observed for *plsC* RBSs. An exception is s5 from cdsA, which has a relatively high frequency considering its TIR of 4,501.

The wider range of RBS strengths for PlsC among the top performers can be interpreted in the light of our previous study reporting that among the intermediate products of the Kennedy pathway only PA accumulated in significant amounts^33^. The conversion of PA by CdsA limits the flux rendering PS production poorly sensitive to the PA production rate and thus also to the amount of PlsC.

### Reverse engineering of clonal variants of the phospholipid synthesis pathway

Due to the repeated PCR steps involved, some mutations accumulated with a frequency >0.04 at off-target locations. To validate the fitness of DNA variants as predicted by the sequencing data, we constructed three clonal variants of expected high fitness with strong RBSs for *plsB*, *cdsA* and *pssA* and varying strengths of the *plsC* RBS (V1-3) (**Fig. 2g (top row), Table S5**), and two clonal variants of expected low fitness, one with weak *cdsA* and *pssA* RBSs (V4) and one with weak *plsB* and *pssA* RBSs and stronger *plsC* RBS (V5). All DNA variants led to PS production in liposomes (**Fig. S13, S14**). However, the population size of the top-producing liposomes differed among the variants. Specifically, we quantified the percentage of liposomes in the strict LactC2-mCherry gate, similar to the sorting gates in the library enrichment phase (i.e., the selection pressure) (**Fig. 2d**). As expected, we found that V1-3 performed well, with 1.4- to 2.2-fold larger gated population sizes (mean values across three replicates) compared to that of the wild type (unpaired *t*-test for WT versus V2 was significant (*P* = 0.05), but those for WT versus V1 or V3 were not). In contrast, V4 and V5 performed poorly. We also found suboptimal performance of a variant with a predicted weak RBS for *plsB* (V6) and good performance of a variant with a predicted strong RBS for *pssA* (V7), as expected from their frequencies in the sorted library as shown in **Fig. 2c**. We also compared the V3, V5 and WT designs on the single-liposome level using confocal fluorescence microscopy. The data suggest improved PS synthesis of V3 over the WT, and decreased activity for V5 (**Fig. S14**). Altogether, the experimentally determined performances of the clonal variants are in agreement with our expectations based on the sequenced libraries.

Next, we quantified the relative protein expression levels for V1-7 using a GreenLys SDS-PAGE assay (**Fig. 2e-g, Fig. S15**) and interpreted the efficiency of the lipid biosynthesis pathway considering these enzyme amounts. This assay confirmed that PS is still produced even with very low PlsC expression (V2), while a reduction in the production of PlsB (V6), or CdsA and PssA (V4), or PlsB and PssA in combination with an increase in PlsC production (V5), reduced PS production significantly (unpaired *t*-tests relative to V2, *P* ≤ 0.05, **Fig. 2d**). High expression of PlsC (V1) did not significantly change PS production compared to low expression (V2). Note that CdsA migrates faster than expected based on its molecular weight (31 kDa), as described earlier^38^.

We found a weak positive linear correlation between the predicted log TIR and the observed protein expression level as quantified from GreenLys SDS-PAGE (R^2^ = 0.3) (**Fig. 2f**). We observed a few instances suggesting that a change in production level of a given protein could influence the production levels of other proteins whose RBSs remained the same (**Fig. 2f**, plots for separate genes). Hence, the same TIR may give different band intensities for different genetic backgrounds. For instance, in V4, upon reducing CdsA and PssA amounts by introducing weak RBSs, the expression levels of PlsB and PlsC increased despite their wild-type RBSs were unmodified (**Fig. S15c**). Furthermore, a *plsC* RBS with TIR 12,360 a.u. led to a higher PlsC production in combination with the downregulation of PlsB and PssA (V5) than in combination with strong RBSs for PlsB, CdsA and PssA (V1) (**Fig. S15c**). On the contrary, a strong reduction in PlsC production in V2 compared to V1 and V3 did not result in a higher expression of the other three proteins. These findings suggest that mutating RBSs of multiple genes leads to unpredictable behavior of the overall expression pattern in PURE system.

### Mutations within the first five codons modulate phospholipid synthesis

We applied a second strategy to modulate the translation initiation rates of the four enzymes of the phospholipid synthesis pathway. The GC content of the sequence directly downstream of the start codon affects the translation rate *in E. coli*, most likely through the tendency of GC-rich sequences to form stronger secondary mRNA structures, which hinder translation initiation^31^. This relationship has recently been observed in PURE system as well^19,39,40^. Inspired by these studies, we varied the GC content of the first six codons between 28-56% for *plsB*, 22-50% for *plsC* and 33-61% for *cdsA* and *pssA*. Only the third base of the codons were mutated such that the mutations were silent, except for the fourth and sixth codons of *cdsA* and *pssA*. A methionine was mutated to isoleucine at the fourth codon of both genes to study a potential positive or negative effect of introducing a second start codon close to the canonical one. At the sixth codon, glycine was mutated to valine to expand the range of GC content, while accepting a potential effect of amino acid substitution.

The theoretical DNA library consisted of 127 (PlsB) × 191 (PlsC) × 127 (CdsA) × 127 (PssA) = 391,241,153 combinatorial (4-site) DNA variants, which is over 33,000-fold larger than the RBS library. Based on the performance of the wild type (**Fig. S9a**), we obtained approximately 3×10^2^-fold screening coverage for the 1-site fraction (572 variants), while only ≤ 1-fold coverage for the fractions with multiple modified target loci (calculation explained in **Table S7**). However, since many DNA variants have the same GC content, we screen the combinatorial GC variants many times (**Table S7**). Therefore, the screening was not an exhaustive search of all combinatorial DNA variants, but of all GC contents.

After four rounds of sorting, we found that some DNA variants were enriched and many variants were depleted from the library (**Fig. S16**). Generally, most DNA variants were enriched in all library fractions, where each library fraction has a different number (*N*) of mutated loci per DNA (*N* = 1, 2, 3 or 4). This observation suggests that these variants were beneficial in different genetic contexts. However, there was no trend visible towards high or low GC contents for the different genes. The most enriched GC variants and their predicted TIRs are listed in **Table S8**.

We also investigated the base-specific effects of each randomized position on PS synthesis (**Fig. 3a**). Site-specific preferences were identified for the tested bases, although no overall preference for A or T over G or C was visible. Interestingly, the effect of the base G at the third position of the second codon was found to be very negative for all genes. This negative effect of the G at this specific position was also discovered in a recent systematic *in vivo* study involving an ultradeep characterization of a large range of randomizations at the 5’-UTR and the start of the coding sequence^31^. Thus, this shows an analogy between *in vivo* and *in vitro* conditions. A few other bases were found to have a severing effect: G in sixth codon of *plsB*, G in fifth codon of *cdsA* and A in second codon of *pssA*. Lastly, changing the second methionine to isoleucine at the fourth codon seems beneficial for both *cdsA* and *pssA*.

**Fig. 3.**
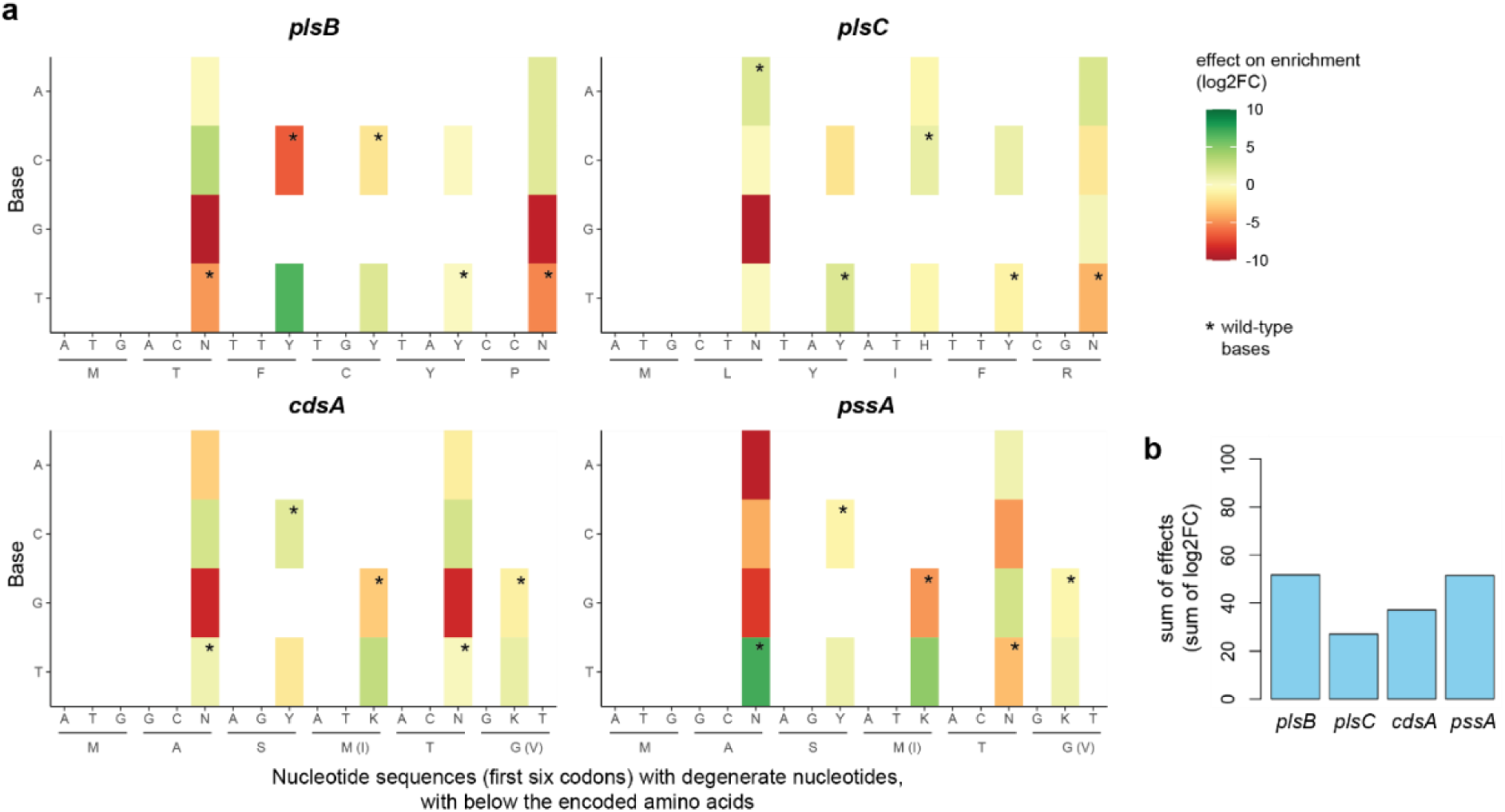
Base-specific effects at specific nucleotide positions in the first six codons of the phospholipid synthesis pathway genes. Data are presented for the 1-site library fraction in R4, but the results across the four library fractions are similar (plots available in the accompanying dataset in the 4TU.ResearchData repository). Number of sequencing reads: 5,827 (*plsB*), 10,254 (*plsC*), 14,594 (*cdsA*), 3,350 (*pssA*). **a** The effect on enrichment (log2FC) was calculated by the fold change of the mean frequency of variants with a certain base at the specific nucleotide position over the mean frequency of variants with the other tested bases. The WT bases are indicated with an asterisk. The encoded amino acids in brackets match with the non-WT bases. **b** Sum of absolute per-base effects on enrichment for each gene.

To determine the total impact of the base substitutions for each gene on PS synthesis, we summed the effects of each tested base. The sum was the highest for *plsB* and *pssA*, suggesting that the mutations in these genes had the largest effect on PS synthesis. In contrast, mutations in the *plsC* gene had the smallest effect on PS production of all genes (**Fig. 3b**). This is in agreement with the RBS library screening which showed that both low and high amounts of PlsC promoted PS synthesis (**Fig. 2d**).

We further investigated the effect of the mutations for the most impactful genes *plsB* and *pssA* by constructing clonal variants with bases yielding a more positive effect for PlsB (V8-11), or a more positive (V10) or negative (V11) effect for PssA (**Table S9**). SDS-PAGE analysis revealed that the PssA expression level was lower with V11, as expected (**Fig. S15c**). However, no significant increase in PlsB or PssA quantity was observed. Nevertheless, a deletion upstream the *plsB* gene of V11 and a deletion upstream the *pssA* gene of V9, which were likely introduced during the cloning of these variants, may have affected the TIR of the proteins.

### Predicting fitness of combinatorial variants based on single variants

Lastly, we examined how predictable the combinatorial variants from the three tested libraries are. Understanding whether combinations of mutations lead to expected or unexpected effects can shed light on the presence of epistatic interactions and clarify whether testing variants in a combinatorial manner is necessary compared to evaluating them individually. Epistasis occurs when the observed fitness (here, variant frequency) deviates from the expected fitness when multiple mutations are combined. For example, the effect of a combination of two RBSs may be larger or smaller than the effect of the first RBS multiplied by the effect of the second RBS (multiplicative model), such that the fitness of the double mutant cannot directly be predicted from analyzing each position in isolation. Epistatic effects may for example appear when multiple genes compete for translation resources. For example, changing the TlR of some genes may indirectly influence the expression level of other genes (improving or severing) by modifying the utilization of tRNAs and tRNA synthetases, a phenomenon that is still poorly understood in cell-free protein synthesis systems.

We first studied the presence of epistatic interactions for the DNA replication RBS library by comparing the observed fitness of the combinatorial variants in the 2-site library fraction with the expected fitness derived from the single variants in the 1-site library fraction. We calculated the expected FC of the combinatorial variants without any epistatic effects by multiplying the FC of each individual RBS (**Fig. 4a**). We then compared the expected and observed sequence landscapes (log2FC, **Fig. 4b**). Both are very similar, with an R^2^ of 0.94, suggesting that no large epistatic interactions govern DNA replication (**Fig. 4c**). Alternative calculations using the 2-site library fraction only (taking the average fold change) or using frequency instead of FC led to similar conclusions (**Fig. S17**). Therefore, the combinatorial variants of this DNA replication library were extremely predictable and, if there was any competition for translation resources, it had minimal effect on DNA self-replication.

**Fig. 4.**
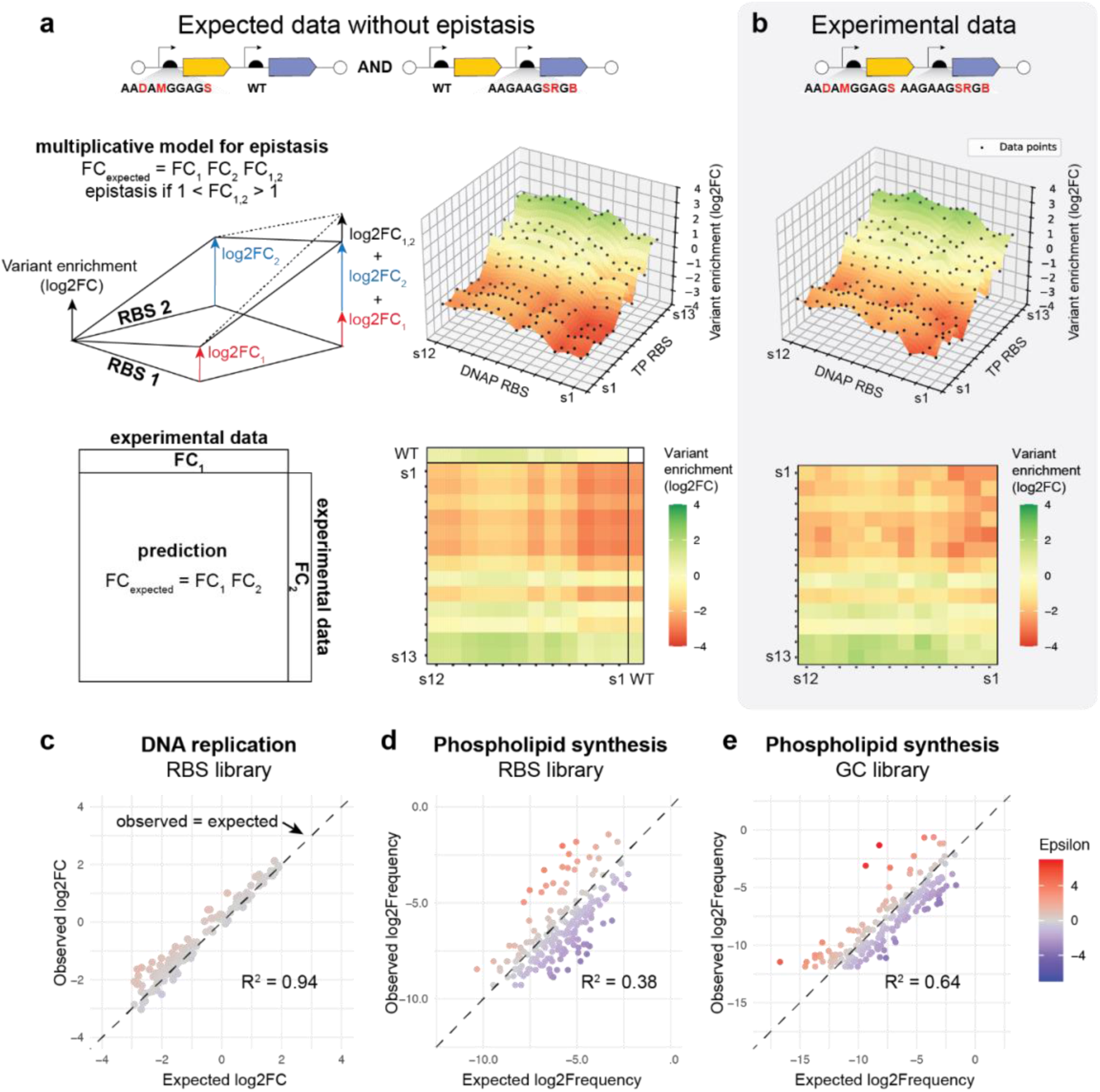
Predicting the fitness of combinatorial variants based on single-mutated variants. **a** Expected combinatorial variant enrichment (log2FC) for the DNA replication RBS library. Prediction was based on a multiplicative model using the enrichment data of DNA variants with one WT and one modified RBS (1-site library fraction). The average of three biological replicates is plotted, where R0 contains 39,928 reads, and R1 33,834 to 56,684 reads per replicate. **b** Experimental data of variant enrichment for the combinatorial library of RBSs of the DNA replicator (average of 3 replicates). **c** Observed versus expected log2FC for the DNA replication RBS library (average of 3 replicates) for all 156 combinatorial variants. **a-c** Data for selection round 1 (R1) were used, because, unlike R2, all DNA variants were detected by Nanopore sequencing, offering a better resolution for low-fitness variants. **d-e** Observed versus expected log2Frequency for the phospholipid synthesis RBS library (d) and GC library (e). Data are from R4 (one biological replicate). Prediction is based on 1-site library fraction. Data points: 205 of the 656 (d) and 189 of the 121,158 (e) pairwise combinatorial variants. **c-e** Dashed line indicates *y* = *x*. Epsilon is the difference between the observed and expected fitness, i.e., Epsilon = log2FC_observed_ – log2FC_expected_ (c), or Epsilon = log2Frequency_observed_ – log2Frequency_expected_ (d-e).

Similarly, we predicted the fitness of the pairwise combinatorial DNA variants of the phospholipid synthesis pathway libraries. Here, we calculated the expected frequency instead of the expected log2FC because of the lower sequencing depth of the pre-sort, 2-site library fraction (24 ± 13 reads per variant for the RBS library) compared to the DNA replication library (186 ± 100 reads per variant). The correlation between the expected versus observed fitness for the two PS synthesis libraries was weaker than that for the DNA replication library (R^2^=0.38 for RBS library, 0.64 for GC library) (**Fig. 4d-e**). Alternative calculations using the 2-site library fraction only or using FC instead of frequency led to similar conclusions (**Fig. S18**). This means that the fitness of these combinatorial variants is less predictable than for the DNA replicators, possibly due to epistatic interactions and/or the more complex sorting workflow, which may increase the level of noise in the sequencing dataset. Nevertheless, it is reasonable that more complex protein systems like the PS synthesis pathway, which involve more genes than the DNA replication system, are more subjected to epistasis affecting gene expression levels as we have also observed from the GreenLys protein gels in **Fig. S15**: RBSs lead to different protein production levels for different genetic backgrounds.

## DISCUSSION

Targeted combinatorial randomization strategies have extensively been applied to optimize expression regulatory sequences for multiprotein systems *in vivo*, for instance genetic circuits^41^ and metabolic pathways^14^, but not in cell-free systems, where the screening of combinatorial libraries is only starting to be explored and so far only targeted single proteins^13,42,43^. Here, we used combinatorial library designs targeting the RBSs or first six codons of coding sequences to tune cell-free expression of multiple genes from multi-gene constructs. Through self-selection or functionality screening, followed by next-generation sequencing (NGS) retrieval of the high-performing variants, we identified variants regulating the functionality of its genetic module at the system’s level. The proposed holistic engineering concept will aid in integrating biological functions in synthetic cells^44,45^.

Several factors may bias the abundance of variants after selection, independent of the mutational effects: the large inherent heterogeneity in functionality within clonal populations of liposomes^46^, differences between variant frequencies in the input library (**Fig. S3a**, **Fig. S12**), and the FACS sorting accuracy of only 80% as determined earlier^47,48^. We assumed that each DNA variant produces an output signal distribution of identical shape (e.g., width of the distribution), leading to a single output quantity that represents its fitness (i.e., amplification fold, or LactC2-mCherry intensity). Increasing the sampling resolution of the variant distributions through implementation of multiple sorting gates could improve the inference of the fitness value and provide insights into the variant’s mechanism of action^49^. In addition, information about low-fitness variants is valuable, especially when combined with active learning-assisted directed evolution. In active learning, machine learning models are used to efficiently navigate large combinatorial sequence spaces, but their predictive accuracy could be improved by including training data on low-fit variants^42,50–52^. Another improvement could be the implementation of unique molecular identifiers (UMIs) during the first 1-2 cycles of PCR to correct for changes in variant frequencies during DNA recovery^53–55^. Despite potential sources of ‘noise’, we demonstrated the functional selection of high-performing variants in one selection round of DNA replication, and within four rounds of sorting for the phospholipid synthesis pathway.

The simultaneous optimization of multiple genes led to mutations that were kilobases apart. Therefore, we used long-read nanopore sequencing for sequence-function mapping. Despite the relatively low single-read per-base sequencing accuracy (99.0-99.7%)^56^, quality control applied to the targeted mutagenesis loci allowed for the detection and counting of DNA variants with minimal mutational distance (i.e., down to one nucleotide difference), as validated earlier^32^. Thus, we linked the mutations within single full-length reads without additional DNA processing steps, such as adding barcodes to identify the DNA variants^43^ or UMIs for deriving consensus sequences from multiple reads^57^.

Our experiments revealed that different target loci have distinct levels of control over the system’s performance. In future optimization campaigns, mutagenesis should preferentially target these most impactful regions to navigate the fitness landscape efficiently, while the expression of other genes could be fixed at a lower rate to save translation resources. We also found that co-expressing multiple proteins with RBSs having theoretical strengths (much) higher than the wild-type RBS, did not lead to large increases in protein production compared to the wild type (**Fig. 1h**, **2f**). Instead, the most significant increases in the amounts of expressed proteins (maximum 1.8 to 2.3-fold relative to the wild type) were obtained in combination with downregulating the co-expression of another protein (**Fig. S15, V4 and V5**), which appears as a more powerful way to balance protein systems and metabolic fluxes than the sole implementation of strong RBSs. More extensive quantitative proteomics analysis would be needed to determine the precise genetic context effects on gene expression and to potentially generate certain design rules that could help optimize expression patterns of multigene constructs in PURE system.

Lastly, pairwise epistasis calculations showed high predictability for the RBS library for self-selection of a DNA replicator and lower predictability for the phospholipid synthesis pathway libraries (**Fig. 4**). These findings underscore the importance of combinatorial scanning over testing each mutation in isolation for different protein systems and/or selection procedures, especially for larger multi-gene systems. However, combinatorial testing comes at the expense of larger library sizes. As mentioned above, active learning strategies can accelerate the search of good designs in a large sequence space.

Other types of libraries could be explored to improve the expression and function of protein systems in synthetic cells. Possible targets include promoters^20,58^, terminators, operator sites, gene order and orientation, promoter insulators (i.e., spacers upstream a promoter sequence)^59^, and operon (multicistronic) designs^20,60^. For example, enhanced transcription can improve protein production^20^, but excess mRNA molecules can sequester initiation factors of the translation machinery, thereby reducing protein synthesis rate^2,61^. Furthermore, protein sequence regions known to govern protein activity, such as active sites or binding sites, could be mutated. Lastly, mutating the type of amino acids in the N- and C-terminal regions, or nucleotide patches that might induce ribosome frameshifting (X/XXY/YYZ) or stalling, might increase the efficiency of protein folding and full-length synthesis^2,61^.

In the future, the system’s level optimization workflow presented here could benefit from the sorting of liposomes based on microscopy images instead of FACS^48^, followed by multi-omics characterization^62–64^. An alternative strategy to variant screening is to couple gene-encoded protein systems to DNA replication, such that a higher fitness of the desired system promotes DNA replication^43,45,65^. In pursuing the development of a synthetic cell, establishing a holistic approach that harnesses combinatorial optimization of multi-protein systems will surely be needed to manage the increasing number of possible genetic designs.

## METHODS

### Reagents and equipment

Chemicals were purchased from Sigma-Aldrich unless stated otherwise. MilliQ water used in buffers and solutions had a resistivity of 18.2 MΩ. DNA concentrations were measured using Qubit dsDNA HS (High Sensitivity) Assay Kit and the Qubit 4 Fluorometer (Invitrogen). Eppendorf Eporator electroporator was used at 1.8 kV. Plasmid isolation was performed with PureYield Plasmid Miniprep System (Promega). PCR-amplified DNA was purified using the QIAquick PCR & Gel Cleanup Kit (Qiagen).

### Biological resources

Bacterial strains *E. coli* K-12 MG1655 and *E. coli* K-12 DH5α were used. pORTMAGE-Ec1 was a gift from the George Church lab (Addgene plasmid #138474; http://n2t.net/addgene:138474; RRID:Addgene_138474). Plasmid G435 was constructed by introducing a point mutation (to remove a PmeI recognition site) in G340, whose synthesis was described earlier^66^. Plasmid G555 (Addgene plasmid #216483; http://n2t.net/addgene:216483; RRID:Addgene_216483) was constructed as described earlier^32^. Plasmids and DNA oligos are listed in **Table S10 and S11**, respectively. LactC2-mCherry was recombinantly expressed and purified as previously described^47^. SSB and DSB were expressed and purified as described in Soengas *et al*., 1995^67^ and Mencía *et al*., 2011^25^, respectively, and stored at 10 mg/mL in 50 mM Tris, pH 7.5, 1 mM EDTA, 7 mM β-mercaptoethanol, 50% glycerol and 60 mM (SSB) or 100 mM (DSB) ammonium sulfate. Purified proteins were aliquoted and stored at −80 °C.

### Design and synthesis of DNA libraries

The RBS libraries for plasmids G435 (encoding DNAP and TP) and G555 (encoding PlsB, PlsC, CdsA and PssA) were designed using the RBS calculator^18^ in the “Optimize Expression Levels” mode (https://salislab.net/software/design_rbs_library_calculator) with parameters: host organism, *E. coli*; target minimum/maximum translation initiation rates, 1/1,000,000; genomic RBS sequence, mRNA sequence from transcription start until (and including) stop codon or terminator. The G555 GC library was designed manually by changing primarily the third bases of the codons to vary GC content of the first six codons of each gene between 22 to 61% (depending on the target locus), while maintaining the originally encoded amino acids (with two exceptions as indicated). The libraries were generated by MOSAIC, a multiplex one-step plasmid diversification protocol, which we recently developed^32^. In this method, DNA oligo libraries are co-electroporated with the target plasmid into *E. coli* cells expressing the single-stranded DNA annealing protein CspRecT from the pORTMAGE-Ec1 plasmid. During plasmid replication in the host cells, the oligos anneal to their target sites and integrate into the plasmid sequence creating a library of plasmids. The (degenerate) DNA oligos used in this study are listed in **Table S11**. For the G435 RBS library, a single electroporation yielded approximately 1,600 colonies, from which the plasmids were isolated. For the G555 RBS library, four electroporation reactions were performed in parallel, yielding in total 3,700 colonies. For the G555 GC library, two parallel electroporation reactions yielded in total 1,800 colonies. The plasmids were isolated from each parallel reaction separately.

PCR was performed on the purified plasmids to linearize the DNA and simultaneously purify the target DNA from the pORTMAGE-Ec1 plasmid. The PCR reaction mix contained 1× Phusion HF buffer, 200 µM dNTPs, 500 nM of each primer (491 ChD and 492 ChD), 2 U Phusion High-Fidelity DNA polymerase and 200 ng plasmid at 100 µL reaction volume. The mixture was split into 2×50 µL, and the thermocycling reaction was performed as follows: 98 °C for 30 s, 20-25 cycles of (98 °C for 5 s and 72 °C for 3 min (G435) or 6 min 20 s (G555)), 72 °C for 5 min. Ten units of DpnI were added to each 50 µL PCR mix and incubated for 1 h at 37 °C to digest the plasmids. The expected PCR products orip2p3 (3,214 bp, derived from G435) and oriPL (6,680 bp, derived from G555) were excised from a 0.7% agarose gel and further purified using QIAquick PCR & Gel Cleanup Kit (Qiagen). The DNA concentration was measured using the Qubit fluorometer. The four oriPL RBS library PCR products were then combined at equimolar amounts to increase library coverage. Similarly, the two oriPL GC library PCR products were combined.

### Synthesis of clonal DNA variants

Clonal plasmid variants were also generated using MOSAIC with a few modifications. Instead of one electroporation, three subsequent rounds of electroporation and plasmid isolation were performed. The *E. coli* strain DH5α harboring pORTMAGE-Ec1 was used instead of MG1655 pORTMAGE-Ec1 to prevent the emergence of smaller plasmids due to recombination during repeated transformation steps. Electroporation was performed with 10-50 ng of plasmid mixture and 2.5 µM of each DNA oligo variant added to the reaction. The plasmids were isolated from colonies grown on ampicillin-supplemented agar plates (50 µg/mL) or from liquid cell culture when the number of colonies on the plates was lower than 50. After three subsequent electroporation steps for mutagenesis, DH5α cells were chemically transformed by heat shock using 3 ng of purified plasmid mixture containing the target plasmids and pORTMAGE-Ec1 to obtain clonally pure plasmids. The transformed cells were plated on ampicillin-supplemented plates to pick single colonies for subsequent plasmid isolation and sequence validation by nanopore sequencing.

### In-liposome gene expression for DNA self-replication

For the expression of DNA libraries in liposomes, swelling solutions were prepared at a volume of 10 µL with 5 µL of PURE*frex* 2.0 (GeneFrontier) Solution I, 0.5 µL of Solution II, 1 µL of Solution III, 0.25 U/µL SUPERase·In RNase inhibitor (Invitrogen), 300 µM of dNTP Mix (Thermo Fisher), 20 mM of ammonium sulfate, 750 µg/mL of purified SSB, 210 µg/mL of purified DSB, and 50 pM of linear orip2p3 DNA. During this study, a change in the commercial PURE*frex* 2.0 solution resulted in impaired DNA replication. Therefore, swelling solutions with clonal DNA variants consisted of an adjusted PURE composition, which yielded better replication compared to the standard kit: 2.5 µL of custom PURE*frex* Solution I lacking tRNAs and NTPs (PFC-Z1608-EX) (i.e., half of the recommended volume), 0.5 µL of Solution II, 1 µL of Solution III, 0.25 U/µL SUPERase·In, 300 µM of dNTP Mix (Thermo Fisher), 20 mM of ammonium sulfate, 750 µg/mL of purified SSB, 210 µg/mL of purified DSB, 50 pM of linear orip2p3 DNA and 1/3^rd^ of the recommended concentrations of NTPs and tRNAs.

The preparation of lipid-coated beads was performed as described earlier^46^ using 50 mol % DOPC, 36 mol % DOPE, 12 mol % DOPG, 2 mol % 18:1 cardiolipin, 0.01 mol % DOPE-Cy5 and 1 mass % DSPE-PEG(2000)-biotin. Five to six milligrams of lipid-coated beads were added to 10 µL of the swelling solution and the tubes were tumbled at 4 °C for 1 h. After four freeze-thaw cycles by dipping the tubes in liquid nitrogen and thawing on ice, 3 µL of liposomes were transferred to PCR tubes containing 0.25 µL of DNase I (2 U/µL) using a cut pipette tip. The liposomes were incubated in a thermocycler at 30 °C for 16 h to enable gene expression, DNA replication, and degradation of non-encapsulated DNA by DNase I. Negative control samples were incubated at 30 °C for 20 min to activate DNase I. The liposomes were then incubated at 75 °C for 15 min to heat-inactive DNase I before amplifying the intraliposomal DNA by (quantitative) PCR.

### Verification of in-liposome DNA replication using quantitative PCR

Quantitative PCR reaction mix was set up at 10 µL volume and contained 5 µL PowerUp SYBR Green Master Mix (Applied Biosystems), 400 nM of each primer (976 ChD and 977 ChD) and 1 µL of heat-inactivated liposome sample diluted 100-fold in MilliQ water. The thermocycling was performed on a Quantstudio 5 Real-Time PCR instrument (Thermo Fisher) as follows: 50 °C for 2 min, 94 °C for 5 min, 45 cycles of (94 °C for 15 s, 56 °C for 15 s and 68 °C for 30 s), 68 °C for 5 min, followed by a melting curve from 65 °C to 95 °C. The DNA concentration was determined using a standard curve of the wild-type DNA with 10-fold dilutions ranging from 0.0001 to 100 pM and the Quantstudio Design and Analysis software v2.6.0 (Thermo Fisher).

### DNA recovery from liposomes after DNA replication

The DNA was recovered from the liposomes by PCR amplification. The PCR reaction mix (100 µL) contained 1× Phusion HF buffer, 200 µM dNTPs, 500 nM of each primer (491 ChD and 492 ChD), 2 U Phusion High-Fidelity DNA polymerase, 5% DMSO and 10 µL of liposome sample diluted 100-fold in MilliQ water. The mixture was split into 2×50 µL, and the thermocycling reaction was performed as follows: 98 °C for 30 s, 30 cycles of (98 °C for 5 s and 72 °C for 3 min), 72 °C for 5 min. The expected PCR products (3214 bp) were excised from a 0.7% agarose gel and further purified using QIAquick PCR & Gel Cleanup Kit (Qiagen). The DNA concentration was measured using a Qubit fluorometer.

### In-liposome gene expression for phospholipid synthesis

The lipid film swelling solutions were prepared at a volume of 20 µL with 10 µL of PURE*frex* 2.0 Solution I, 1 µL of Solution II, 2 µL of Solution III, 0.5 U/µL SUPERase·In, 1 mM CTP, 0.5 mM G3P, 0.5 mM L-serine, 5 mM β-mercaptoethanol, and 10 pM linear oriPL DNA. Lipid-coated beads were prepared and added to the swelling solution as described above, except that the mass of beads was doubled, and the tubes were tumbled at 4 °C for 1 h. After four freeze-thaw cycles, 15 µL of liposomes and 0.5 µL of DNase I (2 U/µL) were transferred to PCR tubes containing a dried film of oleoyl-CoA (Avanti Polar Lipids) to reach a final concentration of 176 µM oleoyl-CoA. The liposomes were incubated in a thermocycler at 37 °C for 6 h (R1), 5 h (R2) or 4 h (R3-5 and clonal variants).

### Liposome screening and sorting with FACS

For each library (RBS or GC library), two identically prepared 15-µL liposome samples were screened and sorted using the BD FACSMelody cell sorter (BD Biosciences). 15 µL of liposomes were diluted in 1,300 µL buffer containing 20 mM HEPES-KOH, pH 7.6, 180 mM potassium glutamate, 14 mM magnesium acetate, and 300 nM LactC2-mCherry. The diluted liposomes were filtered using the 35 µm nylon mesh of a cell strainer cap of 5-mL round bottom polystyrene test tubes (Falcon). LactC2-mCherry fluorescence was measured using a 561-nm laser line and emission filter 610/20 nm. The photon multiplier tube voltages were set at 342 V for FSC, 405 V for SSC (with SSC threshold 359 V), and 461 V for LactC2-mCherry. First, a gate was set on FSC and SSC to select the main liposome population (∼60-80% of all events) (**Fig. S19**). For each sample, 5,000-30,000 liposomes with the highest LactC2-mCherry signal were sorted in purity mode, becoming more stringent in the later sorting rounds. The sorted liposomes from two identical samples were combined. The recorded data was further processed using Cytobank (https://community.cytobank.org/). Here, in addition to the gating based on FSC and SSC, events with LactC2-mCherry signal lower than 2 a.u. were discarded (**Fig. S19**).

### DNA recovery from liposomes sorted by FACS

To recover the DNA from the sorted liposomes, the sorted fractions were spun down at 12,000 g for 3 min, whereafter the supernatant was removed. The remaining ∼10 µL was incubated at 75 °C for 15 min to heat-inactivate DNase I. PCR reactions were prepared at 100-µL reaction volume by mixing 1× Xtreme buffer, 400 µM dNTPs, 300 nM of each primer (1459 ChD and 1460 ChD), 2 units of KOD Xtreme Hotstart DNA polymerase, and 10 µL of heat-inactivated liposome solution. The mixture was split into 2×50 µL, and the thermocycling reaction was performed as follows: 98 °C for 2 min, 35 cycles of (98 °C for 10 s and 68 °C for 6 min 40 s). The expected PCR products (6,616 bp) were excised from a 0.7% agarose gel and further purified using QIAquick PCR & Gel Cleanup Kit (Qiagen). A second PCR reaction was performed with ∼60 ng of the purified DNA using primers 1301 ChD and 1518 ChD to restore the full-length DNA template (6,680 bp), and the amplicon was purified as described above. The DNA concentration was measured using a Qubit fluorometer.

### Nanopore sequencing and analysis

DNA samples were diluted to a concentration of 34 ng/µL (orip2p3) or 71 ng/µL (oriPL). Nanopore sequencing was performed by Plasmidsaurus (Oregon, US). First, analysis of the raw fastq sequencing reads was carried out using the Galaxy web platform^68^ (https://usegalaxy.org). The reads were filtered on length (3,020-3,300 nt for orip2p3, 6,400-7,000 nt for oriPL) using the *Filter FASTQ reads by quality score and length* tool^69^, and mapped to the wild-type sequences using *Map with minimap2*^70^. *Minimap2* was executed using default parameters, except *retain at most INT secondary alignments* that was set to 0. For oriPL data, the *gap open penalty deletion* and *gap open penalty insertion* were set to 16 and 48, respectively.

The mapped reads were further processed using custom R scripts (R v4.4.1, RStudio v2023.09.1+494). A quality control was applied to the mutated regions of interest using a per-base quality cutoff of 50; this quality control method was validated as described earlier^32^. The variant frequency was calculated by dividing the number of reads of a given variant by the total number of reads within the corresponding *N*-site library fraction (i.e., the library fraction containing all sequences with *N* mutated target loci). The variant enrichment (defined as fold change, FC) was calculated by dividing the frequency of a variant by the frequency of that variant in the initial library before sorting (R0). The variant enrichment was plotted as the log2FC (log2 Fold Change). The sum of frequencies per RBS within each *N*-site library fraction, as plotted in **Fig. 2c, S12, S16**, was divided by the number (*N*) of mutated target loci per DNA to normalize the total frequency within each *N*-site library fraction to 1.

### Positional base-specific effects

The positional base-specific effects on variant enrichment were calculated as previously described^31^. The mean frequency of all DNA variants containing a specific base on a specific position was divided by the mean frequency of all DNA variants containing another base on that position. The effect on enrichment was plotted as the log2FC.

### Fitness predictions of combinatorial variants without epistasis

The expected fitness of combinatorial variants was calculated by multiplying the fitness of each individual variant. As the frequency can be seen as the expected probability to sort and sequence a DNA variant, the Bayes’ law about conditional probability can be used to support the choice for the multiplicative model, as suggested by a previous *in vitro* screening study^13^. DNA variants with less than three reads in the 2-site library fraction were excluded from the analysis.

For the DNA replicator library, the expected fitness and the difference between the observed and expected fitness (Epsilon) were calculated as follows:

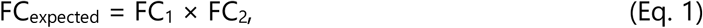

where FC_i_ is the FC of the single variant in the 1-site library fraction.

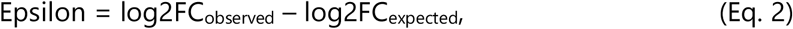

where FC_observed_ is the FC of the combinatorial variant in the 2-site library fraction.

For the PS synthesis libraries, we used frequency instead of FC due to the lower sequencing depth of the pre-sort libraries:

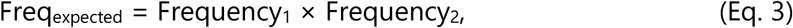

where Frequency_i_ is the frequency of the single variant relative to all variants with that target site mutated in the 1-site library fraction. The difference between the observed and expected fitness is thus defined as:

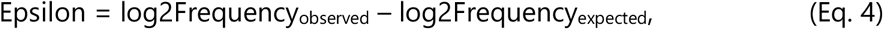

where Frequency_observed_ is the frequency of the combinatorial variant relative to all variants with those two target sites mutated in the 2-site library fraction.

We repeated the calculation of the expected fitness using the 2-site library fraction instead of the 1-site library fraction, and we reached identical conclusions (**Fig. S17-S18**). The expected fitness and Epsilon were calculated for the DNA replicator and PS synthesis libraries as follows:

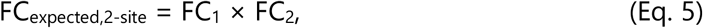

where FC_i_ is the *average* FC of all variants in the 2-site library fraction with mutation *i*.

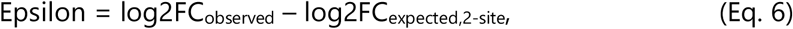

where FC_observed_ is the FC of the combinatorial variant in the 2-site library fraction.

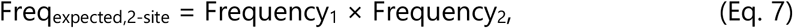

where Frequency_i_ is the *sum* of frequencies of all variants in the 2-site library fraction with mutation *i*.

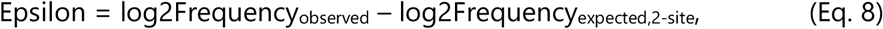

where Frequency_observed_ is the frequency of the combinatorial variant relative to all variants in the 2-site library fraction.

### Off-target mutations

The presence of off-target mutations (outside RBS regions) was analyzed by comparing the mapped reads of the DNA recovered from liposomes with the initial library in the Integrative Genomics Viewer (IGV) v2.16.0 software using a coverage allele-fraction threshold of 0.01 (DNA replication) or 0.04 (PS synthesis libraries).

### GreenLys SDS-PAGE

Relative quantification of protein production was performed by running bulk PURE system reactions. Reactions were set up at 10 µL volume, containing 5 µL of PURE*frex* 2.0 Solution I, 0.5 µL of Solution II, 1 µL of Solution III, 0.25 U/µL SUPERase·In, 0.5 µL of 5-fold diluted FluoroTect Green_Lys_ in vitro Translation Labeling System (Promega) and 1 nM of linear DNA template. The samples were incubated at 30 °C (DNA replication proteins) or 37 °C (phospholipid synthesis enzymes) for 2 h, unless indicated otherwise. Each sample was supplemented with 0.25 µL of RNase A Solution (4 mg/µL) (Promega) and incubated for 30 min at 37 °C. 5 µL of sample was mixed with 1.9 µL of solution containing 1× Laemmli SDS sample buffer, non-reducing (Thermo Fisher), and 10 mM dithiothreitol. The samples were incubated for 5 min at 70 °C to denature the proteins and run on an SDS-PAGE gel consisting of a 12% polyacrylamide resolving gel and a 4% stacking gel. Fluorescently labelled proteins were visualized with a Typhoon fluorescence gel imager (Amersham Biosciences) using a 488-nm laser and a 520-nm band-pass emission filter. Band intensities (adjusted band volume with background subtracted) were quantified using Image Lab v6.1.0 (Bio-Rad Laboratories) relative to the wild-type DNA sample.

### Confocal microscopy and image analysis

Samples for confocal fluorescence microscopy were prepared by mixing 1-2 µL of liposomes with 8 µL of buffer containing 20 mM HEPES-KOH, pH 7.6, 180 mM potassium glutamate, 14 mM magnesium acetate and 5× dsGreen (Lumiprobe). Using a cut pipet tip, the 10-µL sample was transferred to a BSA-functionalized custom-made glass imaging chamber. The sample was incubated at 4 °C for 1-1.5 h before imaging for liposome sinking. Image acquisition was performed with the A1R Laser scanning confocal microscope (Nikon) with an SR Apo TIRF 100× oil immersion objective (NA 1.49) and NIS-Elements v5.30.07 software (Nikon). Imaging was performed using laser line 488 nm with emission filter 525/50 nm (dsGreen), laser line 561 nm with emission filter 595/50 nm (LactC2-mCherry), and laser line 640 nm with emission filter 700/75 nm (DOPE-Cy5). Acquisition settings and adjustments of brightness and contrast were identical for multiple images when comparing fluorescence intensities. Using Fiji (ImageJ2 v2.9.0)^71^, the dsGreen signal inside liposomes was quantified by detecting particles on the binarized dsGreen channel followed by measuring the mean pixel intensity for each particle. The quantification of LactC2-mCherry intensity was performed using SMELDit (https://github.com/DanelonLab/SMELDit).

### Statistics for the number of DNA molecules per liposome

Following Poissonian partitioning, the probability of encapsulating *k* DNA molecules per liposome is:

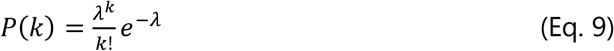

where λ is the expected average number of DNA molecules per liposome, which is calculated as:

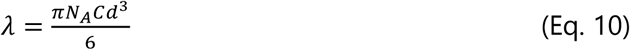

where N_A_ is the Avogadro constant, *C* is the concentration of DNA and *d* is the average liposome diameter.

With 10 pM DNA and a liposome diameter of 4 µm (λ = 0.2), the probability of encapsulating one DNA molecule per liposome is 0.16, and the probability of encapsulating more than one molecule is 0.02. Experimental validation that encapsulation is mostly clonal at 10 pM and 50 pM was performed previously^47^. A more accurate approximation of *λ* values and encapsulation probabilities can be derived by accounting for the experimentally measured distribution of the liposome diameters (https://github.com/DanelonLab/ACS), but the main conclusions remain valid.

### Visualizations

Radar charts, heatmaps and rarefaction plots were generated with R v4.4.1 in RStudio v2023.09.1+494. 3D sequence landscapes were generated with Python in Google Colaboratory. To generate the 3D landscapes, the variants with zero sequencing reads were assigned a log2 fold change identical to the lowest finite number in that dataset. Other graphs were made using Prism v10.0.3 and Illustrator v28.5.

### Statistics

Bar height in bar plots is the mean value across replicates (the individual data points are often appended), and error bars indicate the standard deviation (SD). Box-whisker plots in **Fig. 1f, S5b and S9b** display the median (middle line), the 75/25 percentiles at the boxes and the 90/10 at the whiskers. Statistical tests (unpaired *t*-test, **Fig. 2d, S14**) were performed using GraphPad Prism v10.0.3 and were significant if *P* value (two-tailed) ≤ 0.05. *R*^2^ is the coefficient of determination. The *P* value in linear regression (**Fig. 2f**) indicates whether the slope is significantly different from zero. The number of biological replicates *n* (i.e., separate liposome samples) is stated in the figure captions.

## DATA AVAILABILITY STATEMENT

The data are available in the main manuscript and Supporting Information. Raw data, processed data and the code for data processing underlying this article are available in the 4TU.ResearchData repository with DOI: 10.4121/1d798f6e-7071-4b7d-9d79-7425e4b9d247.

## SUPPORTING INFORMATION

Supplementary figures presenting data of nanopore sequencing, agarose gel electrophoresis, SDS-PAGE, fluorescence microscopy, FACS and qPCR. Supplementary tables listing plasmids, DNA oligos, DNA variant sequences and variant read counts.

## AUTHOR CONTRIBUTIONS

MvdB and CD conceived the project; CD and NJC supervised the project and acquired funding; MvdB and CD designed the experiments; MvdB performed experiments and data analysis and created visualizations; MvdB and CD wrote the paper with input from NJC.

## ACKNOWLEDGEMENTS

We thank Ana Restrepo Sierra for her contribution to the conceptualization and design of the RBS libraries. We are grateful to Ilja Westerlaken for purifying the LactC2-mCherry proteins, Federico Ramirez Gomez for acquiring and analyzing microscopy images of the phospholipid-producing liposomes and Laura Sierra Heras for the adjusted PURE system composition. We also thank Liedewij Laan and Stefan Loonen for fruitful discussions, GeneFrontier for sponsoring our research, and Miguel de Vega and Alicia del Prado from the Centro de Biología Molecular Severo Ochoa (Madrid) for providing purified SSB and DSB proteins. MvdB, CD and NJC acknowledge financial support from the “BaSyC – Building a Synthetic Cell” Gravitation grant (024.003.019) of the Netherlands Ministry of Education, Culture and Science (OCW) and the Netherlands Organisation for Scientific Research (NWO). CD also acknowledges funding from Agence Nationale de la Recherche (ANR-22-CPJ2-0091-01). NJC also acknowledges funding by the NWO-SUMMIT grant “Evolving life from non-life (EVOLF)” (SUMMIT.1.004).

## COMPETING INTERESTS

The authors declare no competing interests.

**Correspondence** and requests for materials should be addressed to Christophe Danelon.

## Notes

### Competing Interest Statement

The authors have declared no competing interest.

