## Supplementary information for "Combinatorial optimization of protein systems in synthetic cells"

**Content:**

Pages S2-16 | Supplementary figures S1-19
Pages S16-24 | Supplementary tables S1-13

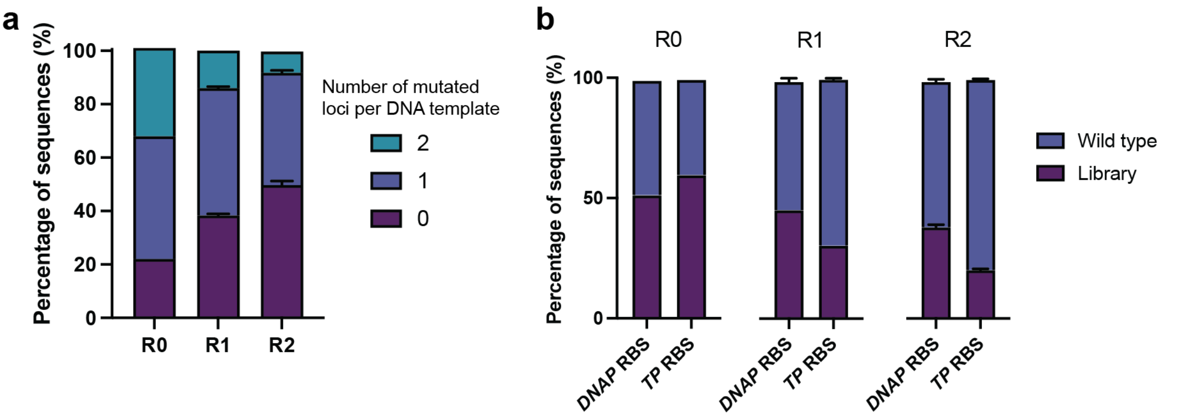


**Fig. S1 Characterization of the orip2p3 RBS library before (R0) and after one (R1) or two (R2) rounds of IVTTR in liposomes. a** Percentage of sequences with zero, one or two modified target loci. Percentage relative to the complete sequenced library after excluding reads with low-quality bases in the target sites. Number of biological replicates: n=1 (R0); n=3 (R1-2). Number of reads: 89,066 (R0); 72,884 (R1, replicate 1); 122,112 (R1, replicate 2); 72,501 (R1, replicate 3); 57,041 (R2, replicate 1); 76,140 (R2, replicate 2); 47,249 (R2, replicate 3). **b** Percentage of wild-type and modified sequences per target locus. The sums of percentages per bar for wild-type and intended library sequences add up to > 98%.

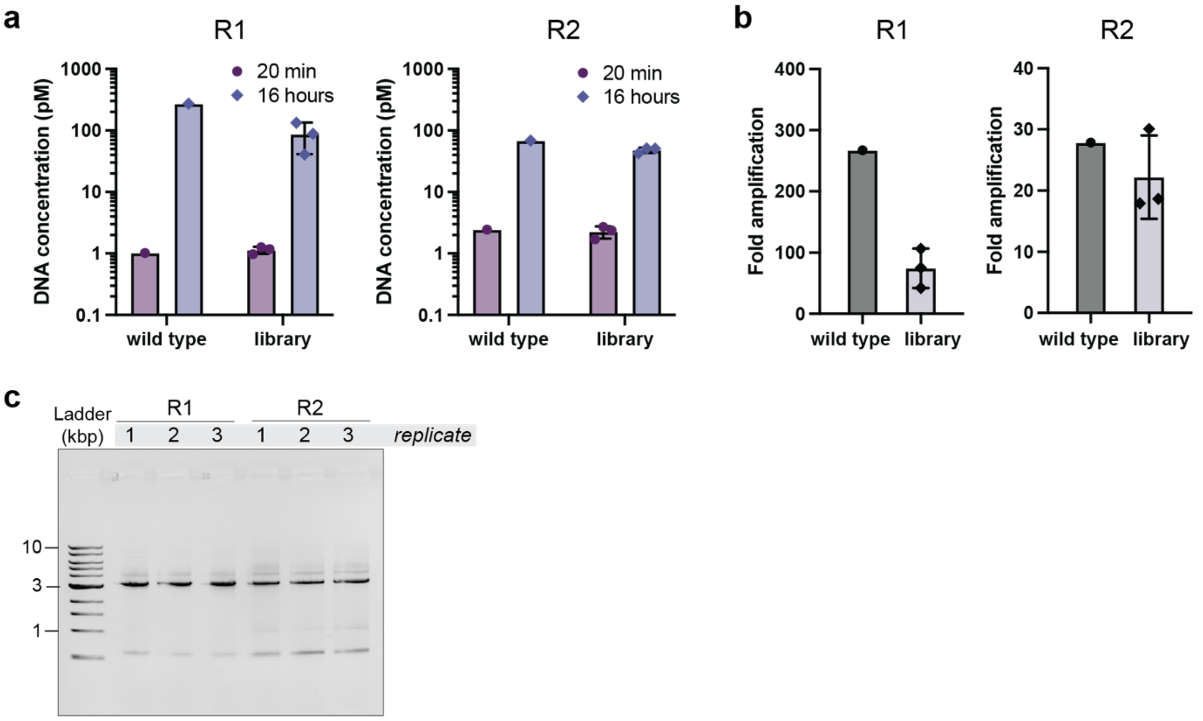


**Fig. S2 Quantification and recovery of wild-type and library orip2p3 templates replicated in liposomes. a** Absolute DNA quantification from in-liposome IVTTR reactions incubated for 16 hours or 20 min (negative control). **b** Replication fold from IVTTR reactions displayed in panel a. Replication fold was determined by the ratio of DNA concentration for reactions incubated for 16 h and 20 min. The wild type sample in round 2 (R2) was the wild-type template recovered from round 1 (R1). **c** PCR-amplified DNA templates (expected size: 3.2 kb) for library IVTTR after 16 h visualized by agarose gel electrophoresis. Purified PCR products were sequenced by nanopore sequencing.

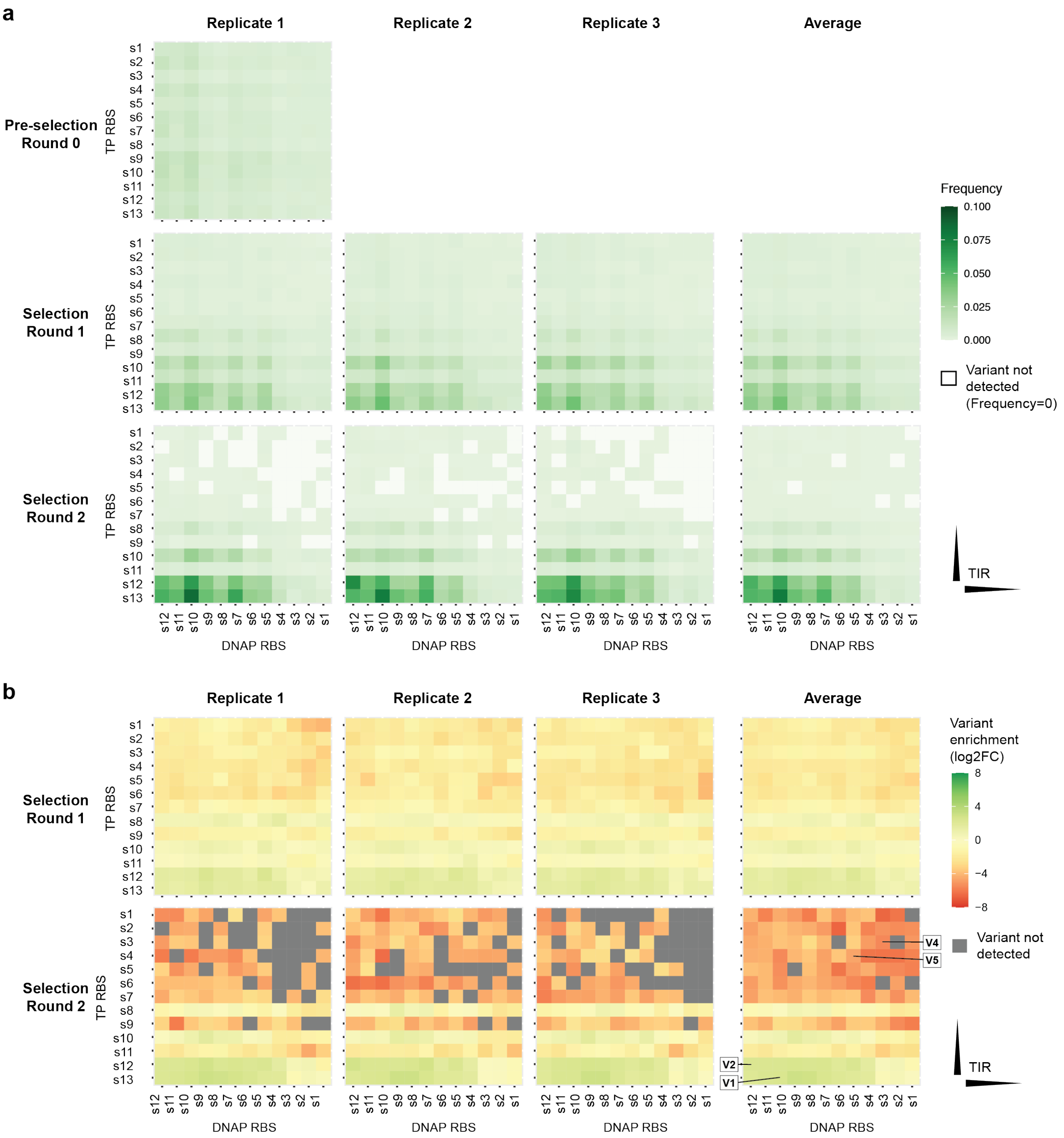


**Fig. S3 Heatmaps displaying the frequency (a) or log2 fold change (b) of variants for each combination of *DNAP* and *TP* RBSs. a** The frequency was calculated by dividing the number of reads by the total number of reads for the fraction of the library containing DNA variants with both RBSs mutated. Number of reads: 28,978 (R0); 10,436 (R1, replicate 1); 16,998 (R1, replicate 2); 10,140 (R1, replicate 3); 4,512 (R2, replicate 1); 6,060 (R2, replicate 2); 3,900 (R2, replicate 3). **b** To calculate the average log2FC, the average frequency was calculated, whereafter this value was divided by the initial frequency of the variant (round 0), and log2 transformed. Clonal DNA variants V1, V2, V4 and V5 are indicated in the right bottom heatmap. In panels a and b the variants are ranked from high to low predicted TIR.

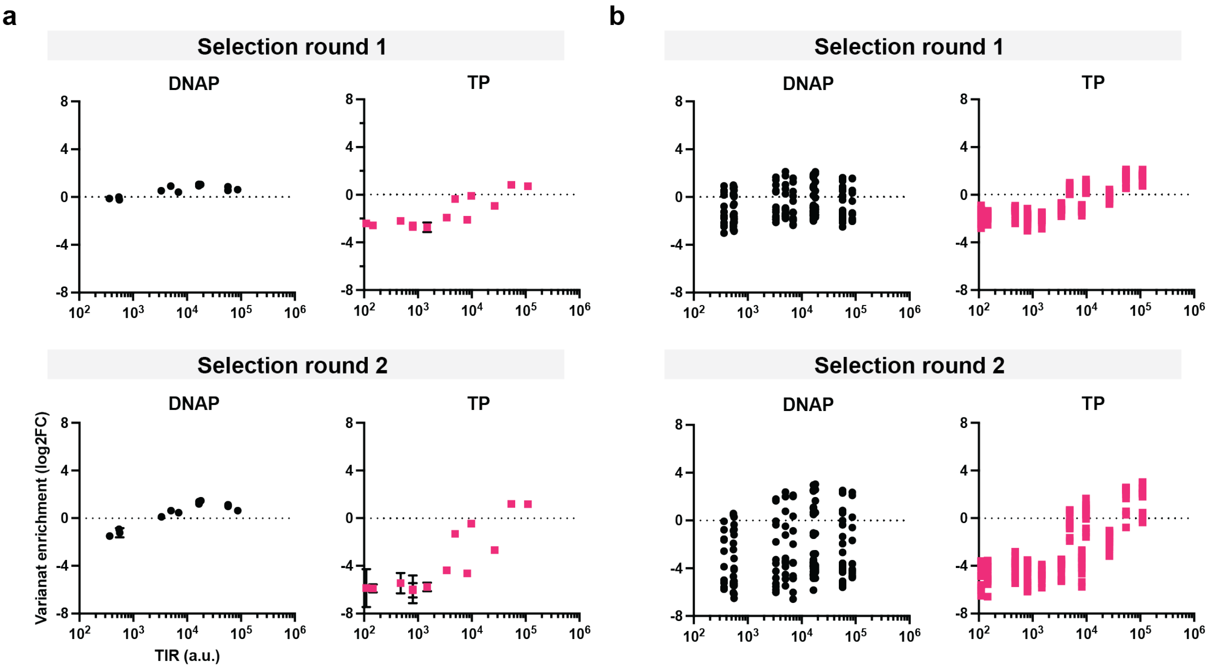


**Fig. S4. Variant enrichment (log2FC) versus predicted TIR for rounds 1 and 2. a** Individual data points for DNA variants that have one wild-type RBS and one mutated RBS. Error bars indicate standard deviation (n=3). **b** Data for the DNA variants that have two mutated RBSs (n=3). For each TIR value, there are multiple data points corresponding to the RBS variants of the other gene on the DNA template.

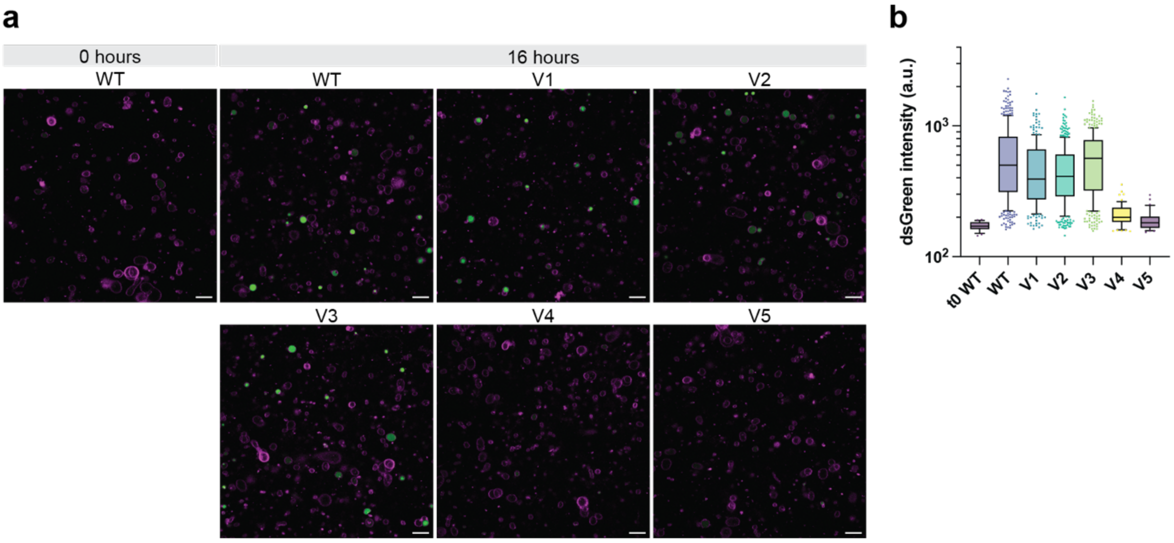


**Fig. S5 Microscopy images of liposomes with replication of clonal DNA variants. a** Additional microscopy images (partly identical to Fig. 1e). Magenta, Cy5-conjugated lipids; green, dsGreen; Scale bars: 10 µm. **b** Second biological replicate for dsGreen quantification in addition to replicate 1 shown in Fig. 1f. Number of dsGreen-positive liposomes: 23 (t0 WT), 364 (WT), 222 (V1), 356 (V2), 340 (V3), 55 (V4), 30 (V5).

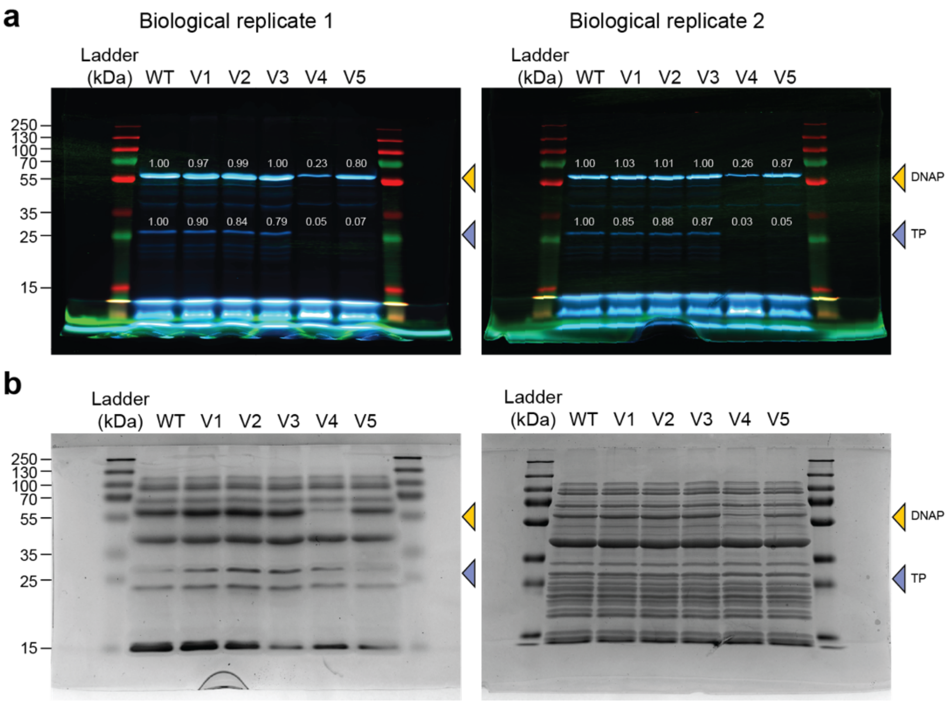


**Fig. S6 Bulk PURE reactions with GreenLys protein labeling for orip2p3 DNA templates with different DNAP and TP RBSs.** Biological replicate 1 was incubated for 8 hours at 30 °C. Biological replicate 2 was incubated for 2 hours at 30 °C. **a** Fluorescence imaging of SDS-PAGE gels for visualization of the bands corresponding to the proteins expressed by PURE system. Band intensities relative to the wild type (WT) are appended in the image. The gel for replicate 2 is also displayed in Fig. 1g. **b** Coomassie staining of the same gels shown in panel a to visualize the bands of all proteins. The staining of the gel for biological replicate 1 was most likely less sensitive due to the use of older Coomassie staining solution.

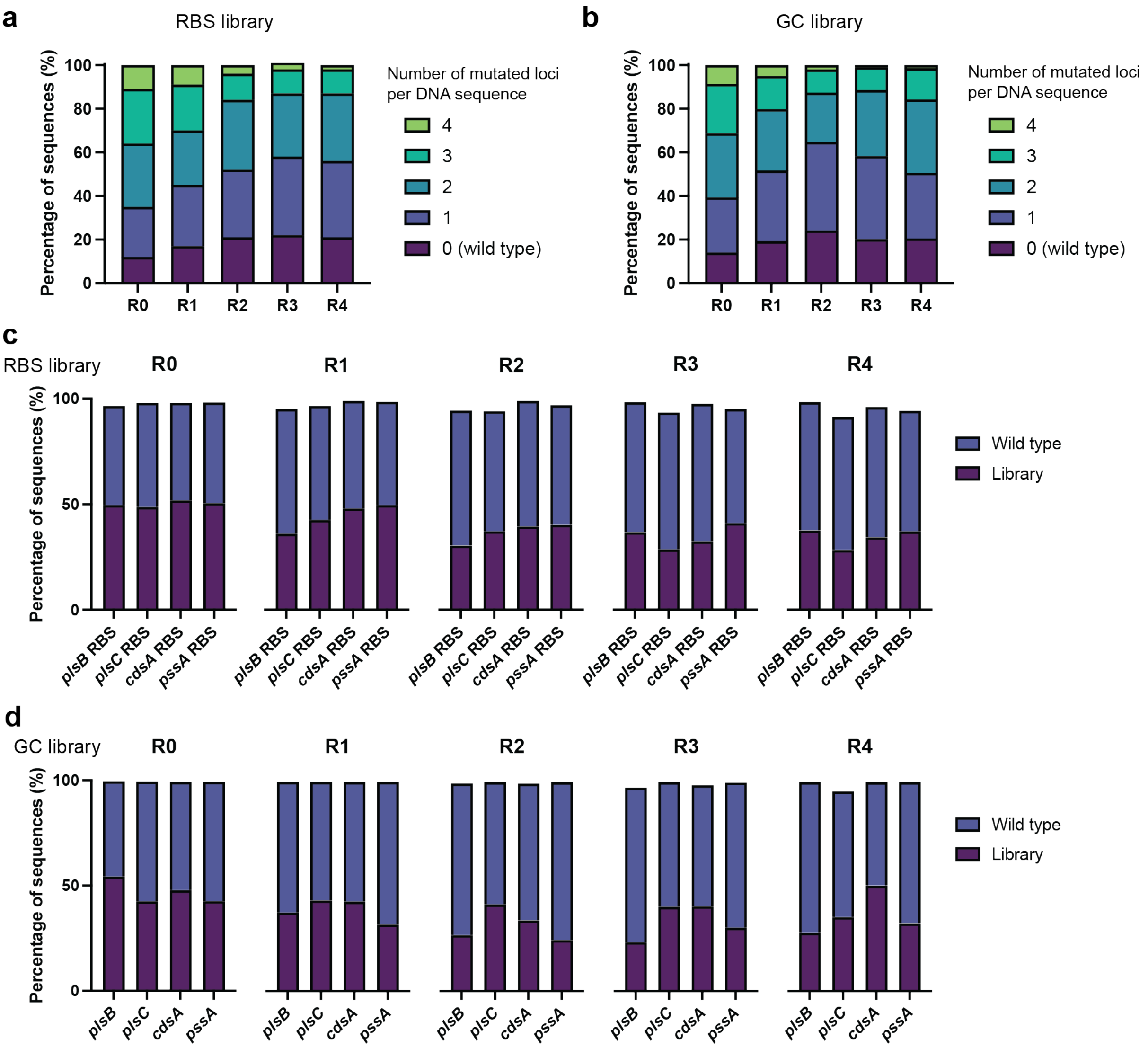


**Fig. S7 Characterization of the phospholipid pathway RBS and GC libraries before (R0) and after one to four (R1-4) rounds of sorting of the liposomes with highest PS production by FACS. a-b** Percentage of sequences with zero, one, two, three and four modified target loci for the RBS library (a) and GC library (b). Percentage relative to the complete sequenced library after excluding reads with low-quality bases in the target sites. Number of reads RBS library (a): 58,988 (R0); 77,727 (R1); 46,248 (R2); 31,776 (R3); 31,777 (R4). Number of reads GC library (b): 41,913 (R0); 116,996 (R1); 142,201 (R2); 118,839 (R3); 120,292 (R4). **c-d** Percentage of wild-type and modified sequences per target locus for the RBS library (c) and GC library (d).

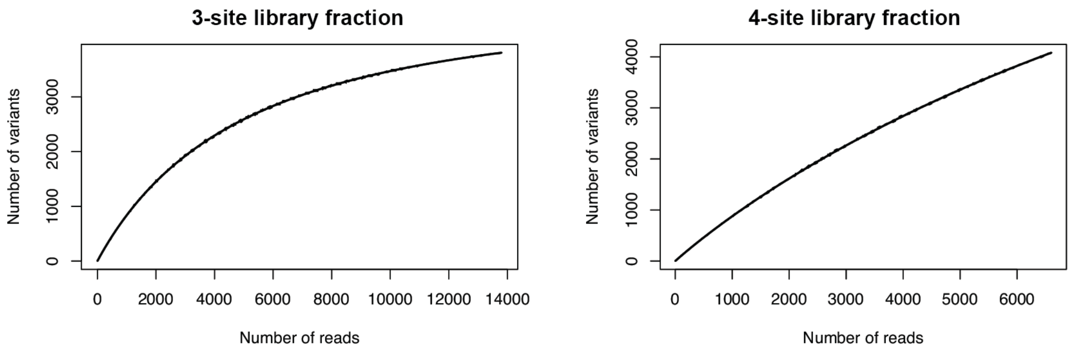


**Fig. S8 Rarefaction plots for DNA variants with three or four RBSs mutated.** Samples were an equimolar mix of four PCR-linearized plasmid library replicates. The graphs do not show a plateau, indicating that the library coverage is most likely larger than the number of variants detected in the nanopore sequencing dataset. For the DNA variants with three mutated RBSs, we detected 3,811 of the 4,518 variants (84%), based on 13,835 reads. For the DNA variants with four mutated RBSs, we detected 4,108 of the 11,583 variants (35%), based on 6,666 reads.

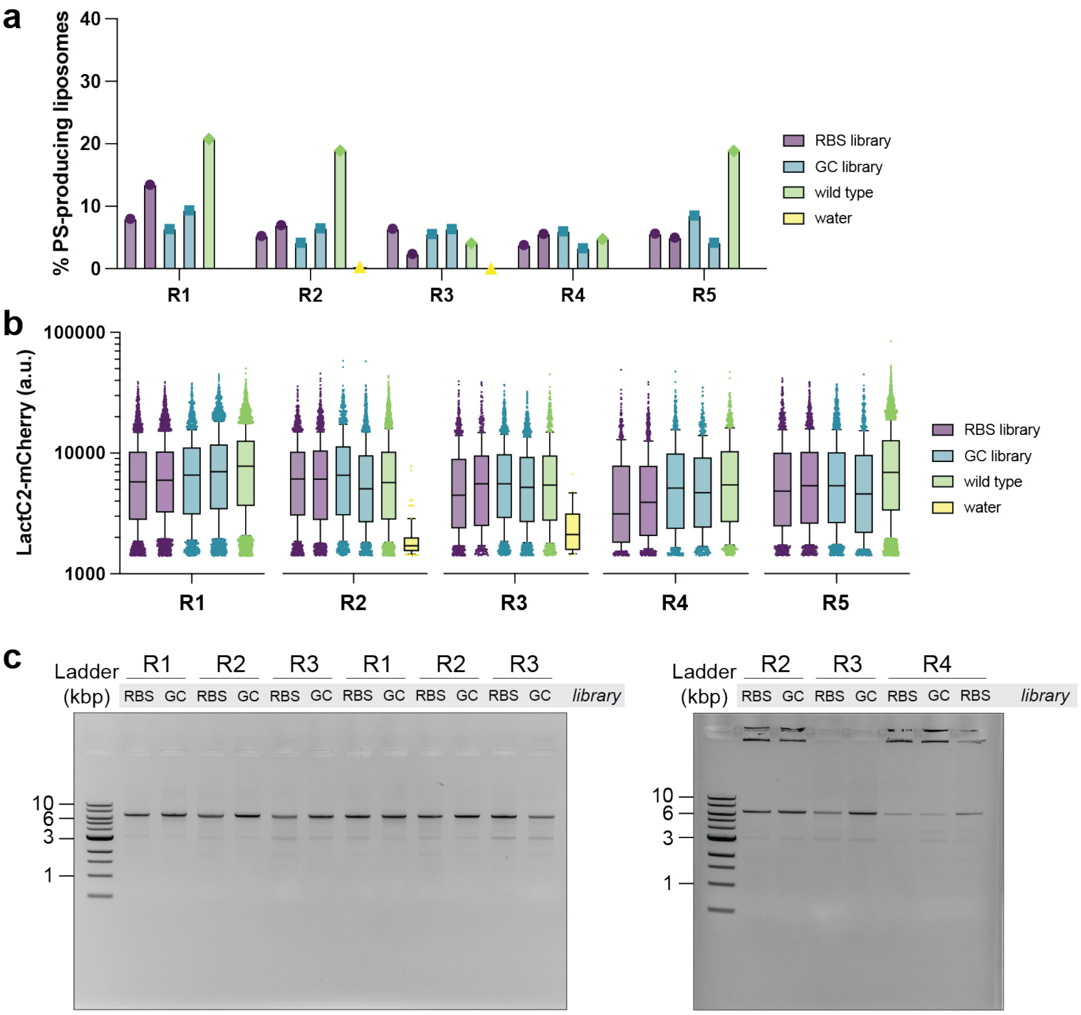


**Fig. S9 FACS sorting and DNA recovery of liposomes producing PS. a** Fraction of PS-producing liposomes measured by FACS. **b** LactC2-mCherry signal of the PS-producing liposomes measured by FACS. **a-b** For the RBS and GC libraries, two biological replicates were analyzed, indicated by identical bar colors. **c** Recovery of DNA from sorted liposomes by PCR (expected template length: 6.7 kb), visualized by agarose gel electrophoresis.

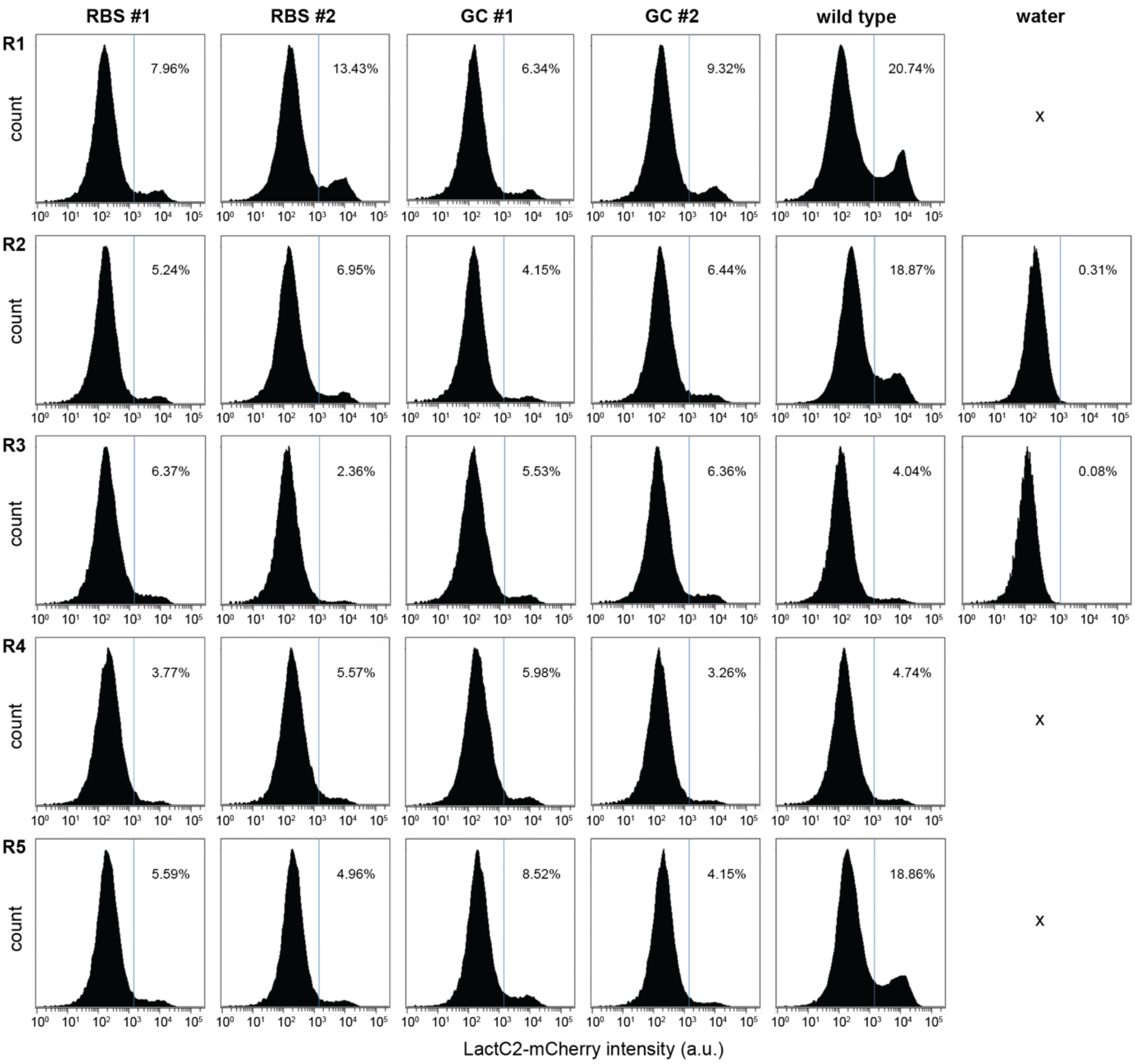


**Fig. S10 Histograms displaying the distribution of LactC2-mCherry intensity of total liposome population.** Liposomes exhibiting an intensity above the threshold, indicated with blue vertical line, are considered PS-producing liposomes. For the RBS and GC libraries, two biological replicates were analyzed, indicated by #1 and #2.

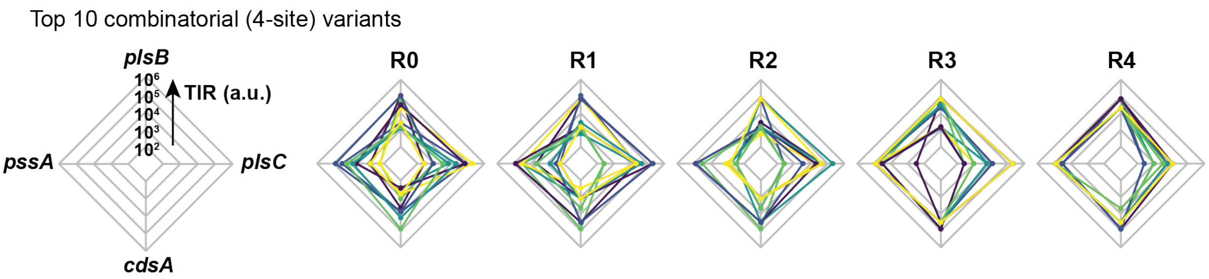


**Fig. S11 Radar charts displaying the top 10 most frequent DNA variants for the phospholipid pathway RBS library for sorting round zero to four.** The top 10 was derived from the sequencing reads containing a library variant for each RBS (i.e., 4-site library fraction).

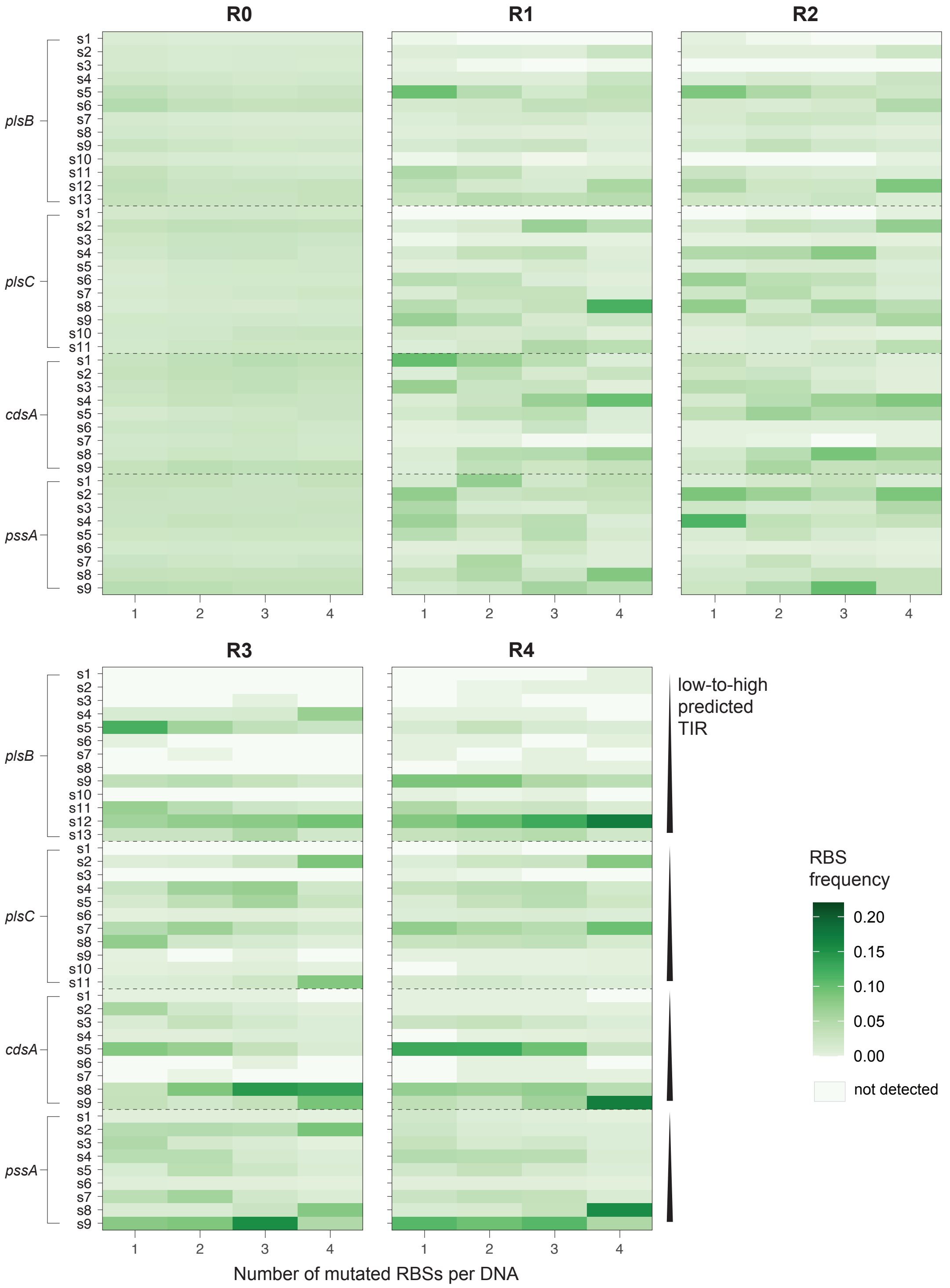


**Fig. S12 Heatmaps showing the frequency of each variant of the PS synthesis RBS library for each round.** The frequency was calculated relative to the library fractions with n number of mutated RBSs per DNA. The heatmap for R4 is also shown in **Fig. 2c**. Number of reads used to calculate the frequency are noted in **Table S12**.

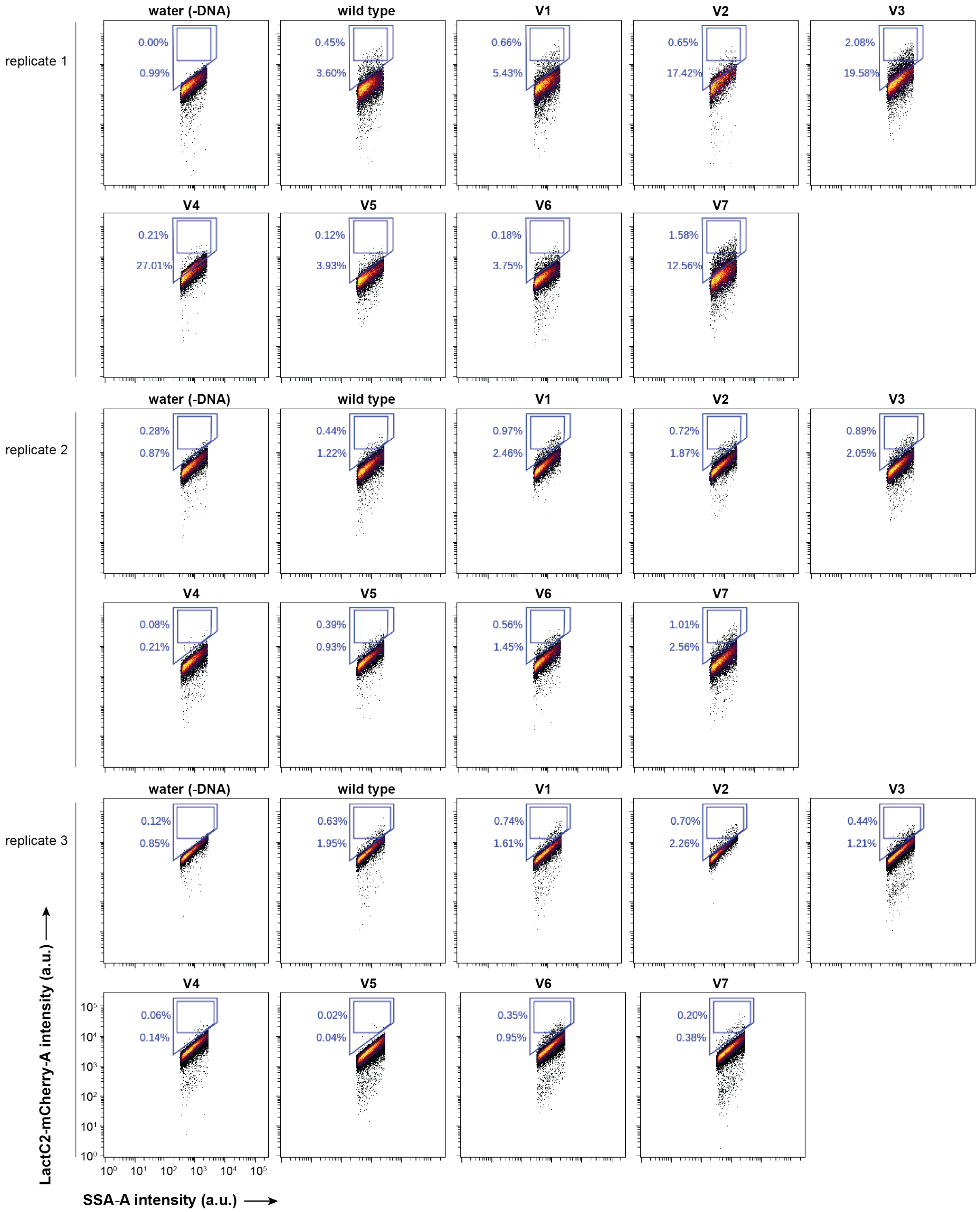


**Fig. S13 Flow cytometry measurements of clonal DNA variants V1-7 for phospholipid synthesis.** 20,000 events per sample (except plot for V2 of replicate 1 which shows 10,000 events).

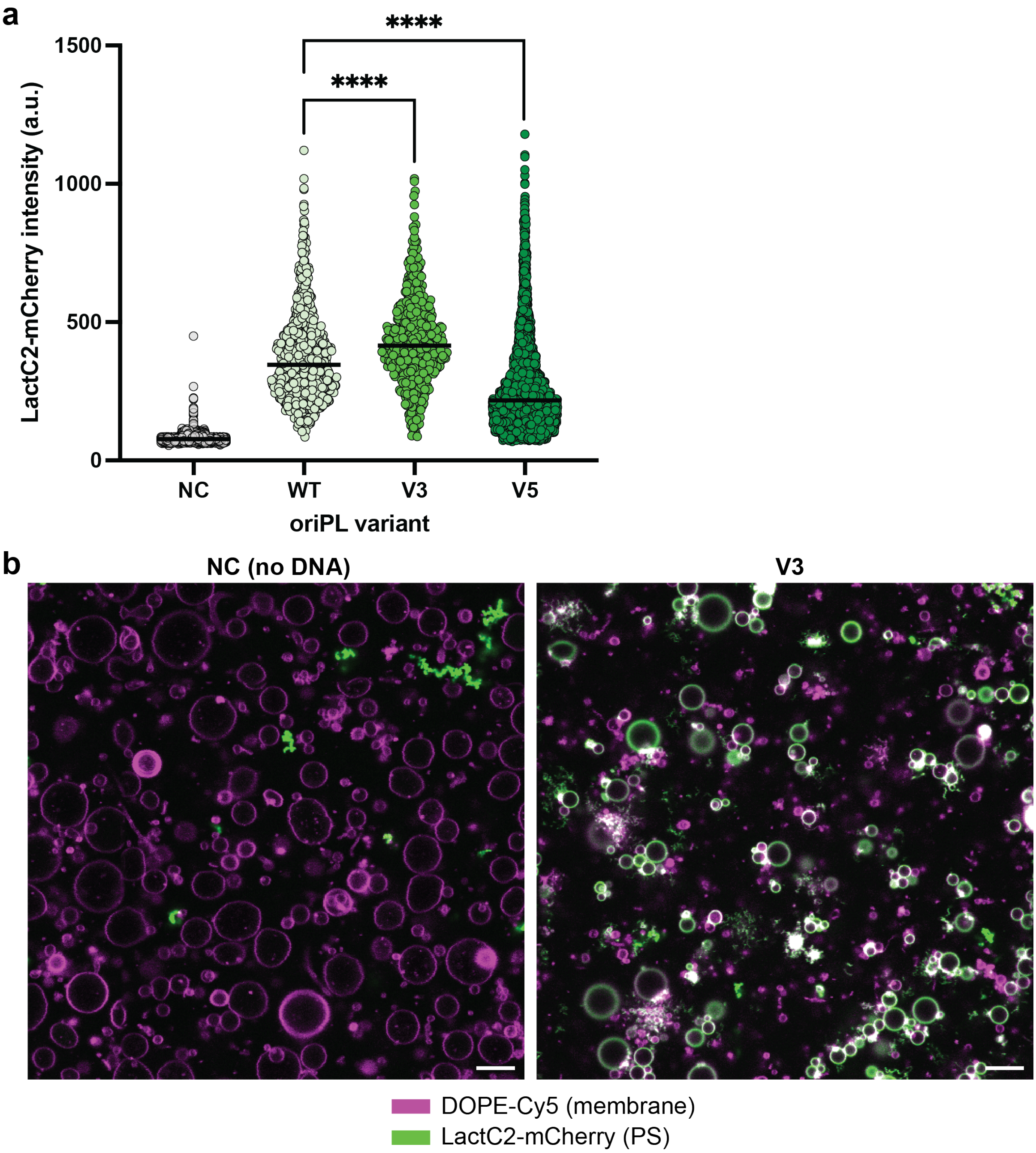


**Fig. S14 Confocal image analysis of LactC2-mCherry fluorescence intensity with different DNA variants.** 1 nM linear DNA was encapsulated in liposomes together with PURE system. The liposomes were incubated for 12 hours at 37 °C. The negative control (NC) contained MilliQ water instead of DNA. **a** Fluorescence intensity profiles from individual liposomes for different samples. Horizontal line indicates the median. Unpaired *t*-test: **** $P < 0.0001$. Analyzed liposomes per sample: 4,188 (NC), 859 (WT), 476 (V3) and 3,566 (V5). **b** Representative confocal fluorescence microscopy images for the negative control (NC) and V3. Scale bars: 10 µm.

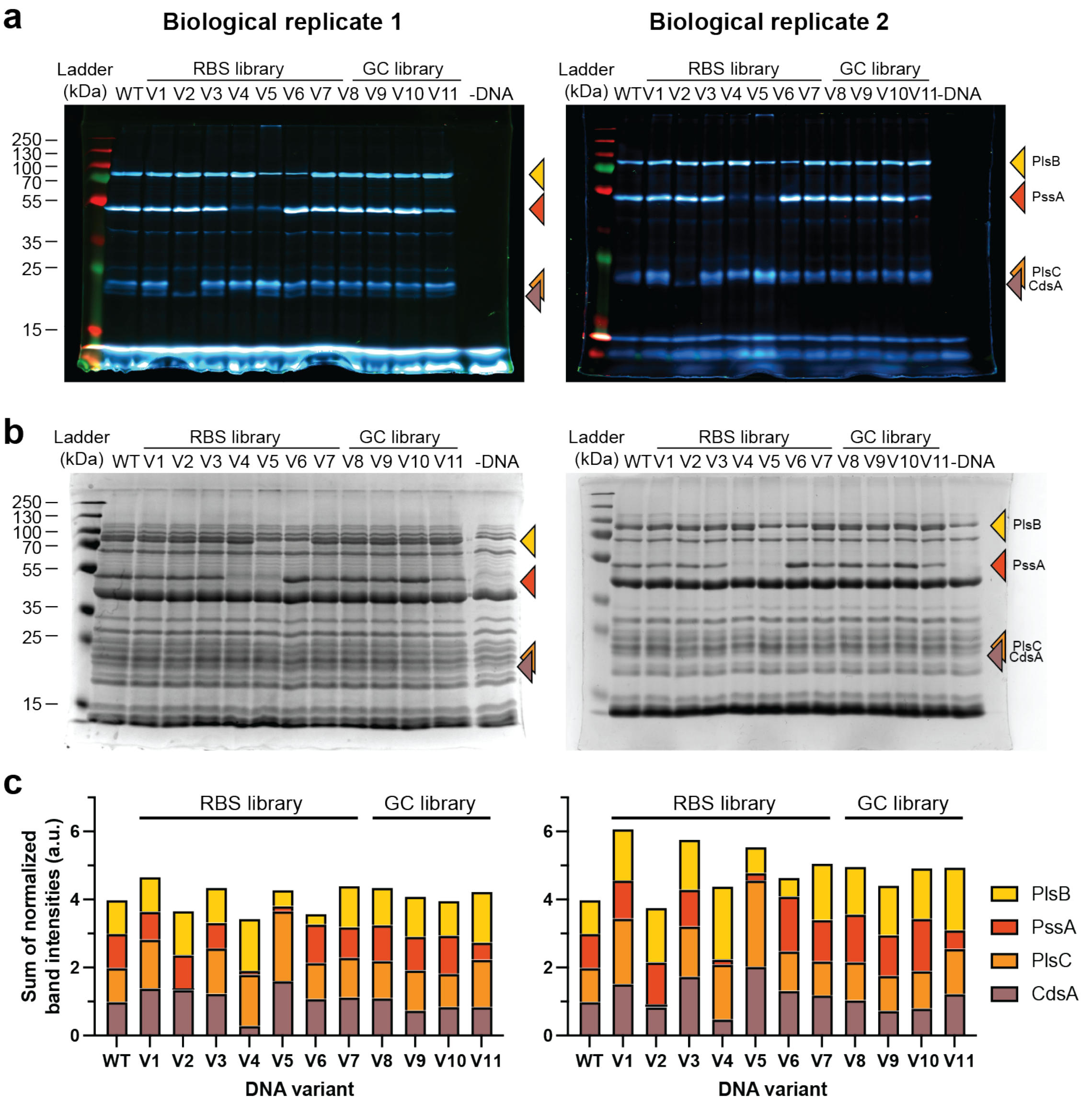


**Fig. S15 Bulk PURE reactions with GreenLys protein labeling for the four phospholipid synthesis enzymes with different RBSs (clonal variants V1-7) or GC content at the start of the coding region (clonal variants V8-11).** The variant sequences and predicted TIRs are listed in Tables S5 and S9. **a** Fluorescence imaging of SDS-PAGE gels for visualization of the bands corresponding to the proteins expressed by PURE system. The gel for replicate 1 is also displayed in Fig. 2e. **b** Coomassie staining of the same gels shown in panel a to visualize the bands of all proteins. **c** Quantification of the band intensities relative to the wild type (WT). Note that V9 and V11 have 4 (unintended) deletions upstream the *pssA* gene, and 1 deletion upstream the *plsB* gene, respectively, which may bias the interpretation of the effect of the targeted GC mutations for these variants.

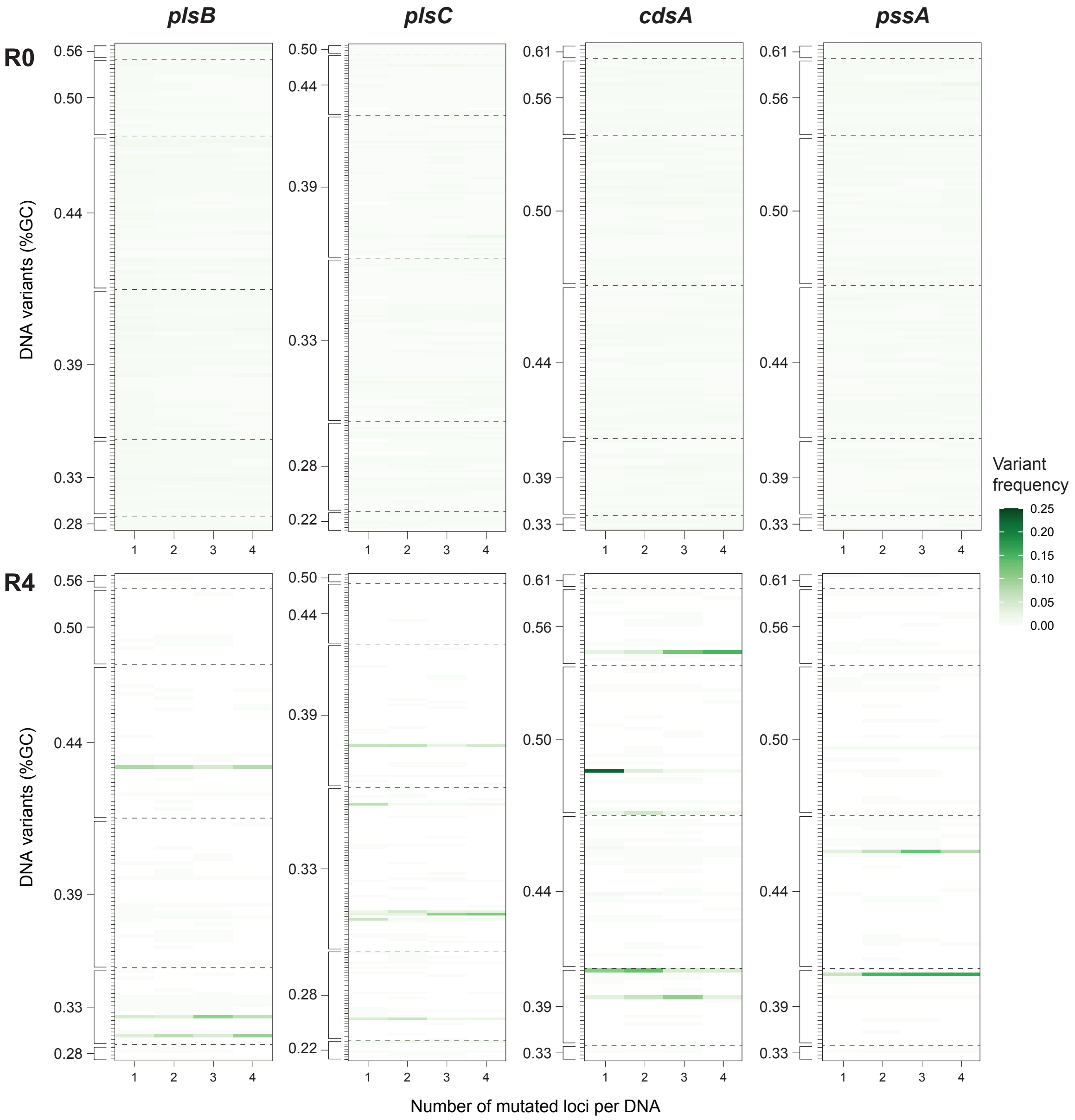


**Fig. S16 Heatmaps showing the frequency of each DNA variant in the PS synthesis GC library for R0 (top row) and R4 (bottom row).** The frequency was calculated relative to the library fractions with n number of mutated RBSs per DNA. Each tick mark on the y-axis corresponds to one DNA variant (127 DNA variants for *plsB*, *cdsA* and *pssA*, and 191 variants for *plsC*). The DNA variants are grouped based on the GC content of the first six codons of the genes, which ranged from 28-56% for *plsB*, 22-50% for *plsC* and 33-61% for *cdsA* and *pssA*. The GC content in the first six codons of wild-type *plsB*, *plsC*, *cdsA*, *pssA* are 39%, 28%, 50% and 50%, respectively. The number of reads used to calculate the frequency are noted in **Table S13**.

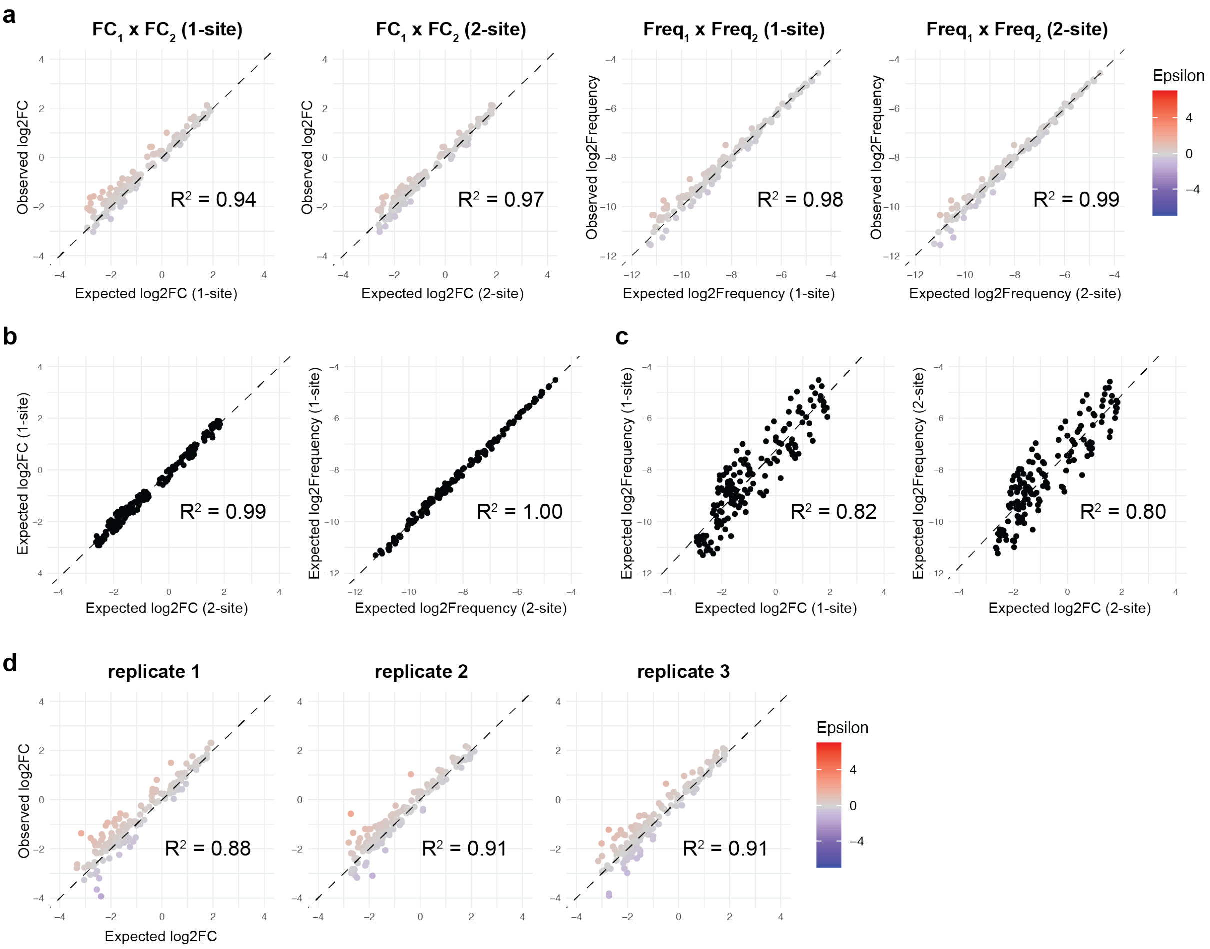


**Fig. S17 Different methods for calculating the expected fitness of combinatorial DNA variants for the DNA replicator RBS library.** Data of selection round 1 (R1). Dashed line is y = x (panels **a** and **d**) or the result of linear regression (panels **b-c**). All plots contain 156 data points. **a** Observed versus expected fitness plots, where fitness is log2FC or log2Frequency, and the expected fitness was calculated using the 1-site or 2-site library fraction data. Epsilon is the log2 observed fitness – log2 expected fitness. **b** The use of the 1-site or 2-site library fraction to calculate the expected fitness has similar outcomes. **c** Correlation between frequency and fold change for calculating the expected variant fitness. **a-c** Average of three biological replicates. **d** Observed versus expected log2FC of the three individual biological replicates. The expected log2FC was calculated using the 1-site library fraction data.

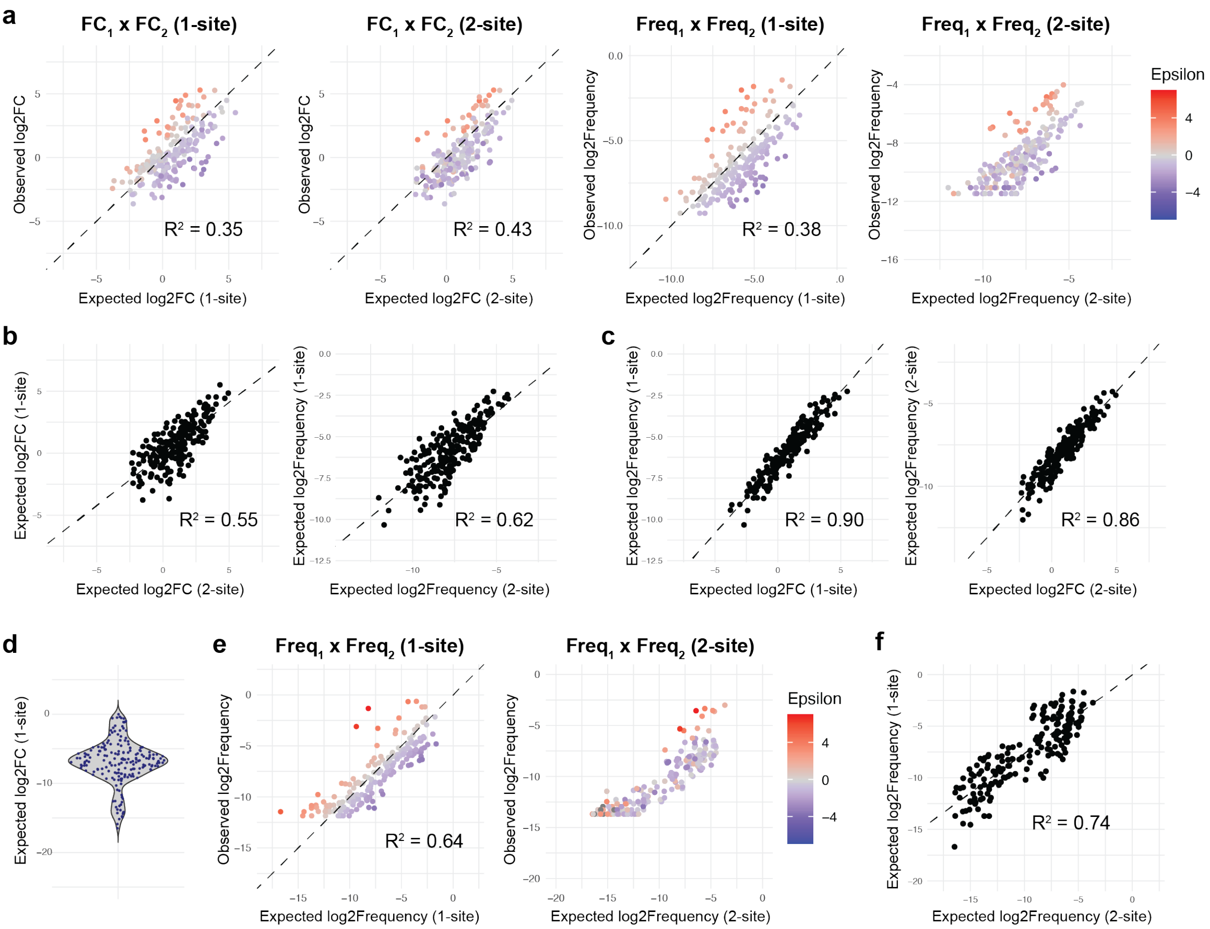


**Fig. S18 Different methods for calculating the expected fitness of pairwise combinatorial DNA variants for the PS synthesis RBS and GC libraries.** Data of sorting round 4 (R4). Dashed line is y = x (panels **a**, **e**) or the result of linear regression (panels **b-c**, **f**). **a-d** RBS library. **e-f** GC library. **a** Observed versus expected fitness plots for the RBS library, where fitness is log2FC or log2Frequency, and the expected fitness was calculated using the 1-site or 2-site library fraction data. Note that expected log2Frequency (2-site) does not scale with observed log2Frequency in the rightmost plot, because the full combinatorial library (with 4 mutagenic sites) is larger than the pairwise variants. The plotted Epsilon is identical for all plots to show that the high-epistasis variants (in red) are detected by all calculation methods (Epsilon is the observed log2Frequency – expected log2Frequency (1-site) for all plots). **b** Correlation between the expected fitness calculated using the 1-site versus 2-site fraction of the RBS library. **c** The use of FC or Frequency as a measure of the variant fitness leads to similar results. **a-c** All plots show 205 of the 656 pairwise combinatorial variants (DNA variants with less than three reads in the 2-site library fraction were excluded from the analysis). **d** Expected log2FC (calculated using the 1-site library fraction) for combinatorial DNA variants with less than three reads in the 2-site library fraction. The expected log2 fold change was low (mostly negative enrichment factors) compared to the sequenced combinatorial variants in panel a (left plot), explaining why the combinatorial DNA variants were not sequenced. 200 variants were plotted. **e** Observed versus expected fitness plots for the GC library. Note that expected log2Frequency (2-site) does not scale with observed log2Frequency in the right plot, because the full combinatorial library (with 4 mutagenic sites) is larger than the pairwise variants. The plotted Epsilon is identical for the two plots to show that the high-epistasis variants (in red) are detected by the two calculation methods (Epsilon is the observed log2Frequency –expected log2Frequency (1-site)). The left plot is identical to Fig. 4e. **f** Correlation between the expected fitness calculated using the 1-site versus 2-site fraction of the GC library. **e-f** 189 of the 121,158 pairwise combinatorial variants are plotted.

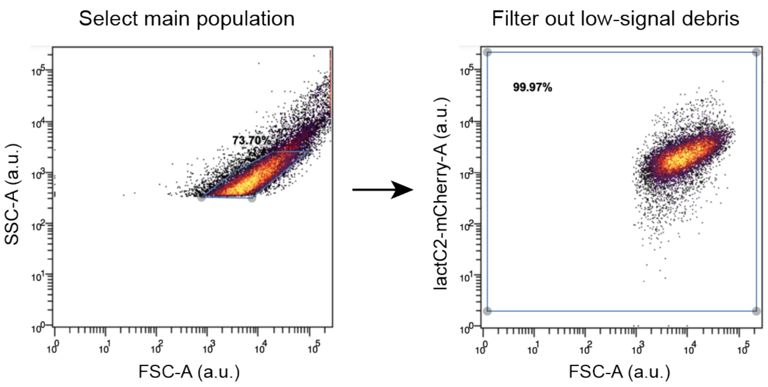


**Fig. S19 Example of gating strategy for flow cytometry data.** Only the first step (gating based on forward and side scatter light) was applied during sorting. The two filtering steps were applied during post-measurement analysis.

**Table S1 List of RBS sequences for DNAP and TP in orip2p3 DNA template.** Red color indicates degenerate nucleotides.

| | DNAP RBS | | TP RBS | |
|---|---|---|---|---|
| | **Sequence** | **Predicted TIR** | **Sequence** | **Predicted TIR** |
| Wild type | TTTAAGAAGGAGATATACAT | 9,123 | TTTAAGAAGGAGATATACTA | 25,228 |
| Library degenerate sequence | TTTAADAMGGAGSTATACAT | - | TTTAAGAAGSRGBTTGACTA | - |
| s1 | TTTAAAACGGAGCTATACAT | 362 | TTTAAGAAGCGGCTTGACTA | 110 |
| s2 | TTTAAGACGGAGCTATACAT | 543 | TTTAAGAAGCGGTTTGACTA | 146 |
| s3 | TTTAATACGGAGCTATACAT | 555 | TTTAAGAAGCGGGTTGACTA | 477 |
| s4 | TTTAAAAAGGAGCTATACAT | 3,329 | TTTAAGAAGCAGTTTGACTA | 801 |
| s5 | TTTAAGAAGGAGCTATACAT | 4,992 | TTTAAGAAGCAGCTTGACTA | 801 |
| s6 | TTTAATAAGGAGCTATACAT | 6,933 | TTTAAGAAGCAGGTTGACTA | 1,476 |
| s7 | TTTAAGACGGAGGTATACAT | 16,525 | TTTAAGAAGGGGCTTGACTA | 3,425 |
| s8 | TTTAAAACGGAGGTATACAT | 16,525 | TTTAAGAAGGAGCTTGACTA | 4,909 |
| s9 | TTTAATACGGAGGTATACAT | 17,920 | TTTAAGAAGGGGTTTGACTA | 8,224 |
| s10 | TTTAAGAAGGAGGTATACAT | 57,485 | TTTAAGAAGGAGTTTGACTA | 9,818 |
| s11 | TTTAAAAAGGAGGTATACAT | 57,485 | TTTAAGAAGGGGGTTGACTA | 26,545 |
| s12 | TTTAATAAGGAGGTATACAT | 86,579 | TTTAAGAAGGAGGTTGACTA | 54,294 |
| s13 | - | - | TTTAAGAAGGAGGTTTACTA | 110,313 |

**Table S2 Clonal DNA variants from orip2p3 RBS library.** The *plsC* RBS of V3 (with predicted TIR: 938 a.u.) was not a variant from the theoretical library, but likely originated from a mutation during MOSAIC or PCR.

| | **DNAP RBS** | | **TP RBS** | |
|---|---|---|---|---|
| | **Sequence** | **Predicted TIR** | **Sequence** | **Predicted TIR** |
| WT | WT | 9,123 | WT | 25,228 |
| V1 | s10 | 57,485 | s13 | 110,313 |
| V2 | s12 | 86,579 | s12 | 54,294 |
| V3 | TTTAATAAGGAGATATACAT | 14,117 | s13 | 110,313 |
| V4 | s3 | 555 | s3 | 477 |
| V5 | s5 | 4,992 | s4 | 801 |

**Table S3 List of RBS sequences for *plsB*, *plsC*, *cdsA* and *pssA* in oriPL DNA template.** Red color indicates degenerate nucleotides.

| | ***plsB* RBS** | | ***plsC* RBS** | |
|---|---|---|---|---|
| | **Sequence** | **Predicted TIR** | **Sequence** | **Predicted TIR** |
| Wild type | TTTAAGAAGGAGATATACAT | 5,896 | TTTAAGAAGGAGATATACAT | 4,947 |
| Library degenerate sequence | TTTAAGGAGGKAAAMBCCAT | - | TTTAAGGVGAATADATACAT | - |
| s1 | TTTAAGGAGGGAAACCCCAT | 246 | TTTAAGGCGAATAGATACAT | 144 |
| s2 | TTTAAGGAGGGAAACTCCAT | 608 | TTTAAGGCGAATATATACAT | 249 |
| s3 | TTTAAGGAGGGAAACGCCAT | 981 | TTTAAGGCGAATAAATACAT | 878 |
| s4 | TTTAAGGAGGGAAAACCCAT | 1,309 | TTTAAGGGGAATAGATACAT | 3,327 |
| s5 | TTTAAGGAGGGAAAAGCCAT | 1,561 | TTTAAGGGGAATATATACAT | 5,725 |
| s6 | TTTAAGGAGGGAAAATCCAT | 2,803 | TTTAAGGAGAATAGATACAT | 7,182 |
| s7 | TTTAAGGAGGTAAACCCCAT | 6,346 | TTTAAGGAGAATATATACAT | 12,360 |
| s8 | TTTAAGGAGGTAAACTCCAT | 15,661 | TTTAAGGGGAATAAATACAT | 20,220 |
| s9 | TTTAAGGAGGTAAAACCCAT | 21,409 | TTTAAGGAGAATAAATACAT | 43,653 |
| s10 | TTTAAGGAGGTAAACGCCAT | 25,289 | TTTAAGGAGGTAAGATACAT | 68,987 |
| s11 | TTTAAGGAGGTAAAAGCCAT | 31,839 | TTTAAGGAGGTAATATACAT | 201,909 |
| s12 | TTTAAGGAGGTAAAATCCAT | 72,247 | | |
| s13 | TTTAAGGAGGTAATATACAT | 112,986 | | |

| | ***cdsA* and *pssA* RBSs** | |
|---|---|---|
| | **Sequence** | **Predicted TIR** |
| Wild type | TTTAAGAAGGAGATATACAT | 5,198 |
| Library degenerate sequence | TTTAAGARKGAGKTATACAT | - |
| s1 | TTTAAGAATGAGTTATACAT | 174 |
| s2 | TTTAAGAGTGAGTTATACAT | 289 |
| s3 | TTTAAGAGGGAGTTATACAT | 659 |
| s4 | TTTAAGAAGGAGTTATACAT | 1,231 |
| s5 | TTTAAGAATGAGGTATACAT | 4,501 |
| s6 | TTTAAGAGTGAGGTATACAT | 7,484 |
| s7 | TTTAAGAGGGAGGTATACAT | 17,054 |
| s8 | TTTAAGAAGGAGGTATACAT | 31,879 |
| s9 | GATAAGGAGGTGATATACAT | 78,824 |

**Table S4 Calculation of FACS screening coverage for RBS library.** Calculation based on 4,500,000 screened events and 13% PS-producing liposomes (for wild type). Calculation example (row 2): Per-variant screening coverage = 4,500,000 (number of events) * 0.23 (frequency of this library fraction of total library) * 0.13 (% PS-producing liposomes) / 42 (number of variants) = 3,204.

| **Library fraction** | **Number of mutated RBSs in this library fraction** | **Number of variants** | **Fraction of total library** | **Per-variant screening coverage** |
|---|---|---|---|---|
| 1 | 0 (WT) | 1 | 0.12 | 70,200 |
| 2 | 1 | 42 | 0.23 | 3,204 |
| 3 | 2 | 656 | 0.29 | 259 |
| 4 | 3 | 4,518 | 0.25 | 32 |
| 5 | 4 | 11,583 | 0.11 | 6 |

**Table S5 Clonal DNA variants from oriPL RBS library.**

| | ***plsB* RBS** | | ***plsC* RBS** | | ***cdsA* RBS** | | ***pssA* RBS** | |
|---|---|---|---|---|---|---|---|---|
| | **Sequence** | **Predicted TIR** | **Sequence** | **Predicted TIR** | **Sequence** | **Predicted TIR** | **Sequence** | **Predicted TIR** |
| WT | WT | 5,896 | WT | 4,947 | WT | 5,198 | WT | 5,198 |
| V1 | s12 | 72,247 | s7 | 12,360 | s9 | 78,824 | s9 | 78,824 |
| V2 | s12 | 72,247 | TTTAAGG GGAATAG ATATAT | 938 | s9 | 78,824 | s9 | 78,824 |
| V3 | s12 | 72,247 | s11 | 201,909 | s9 | 78,824 | s9 | 78,824 |
| V4 | WT | 5,896 | WT | 4,947 | s1 | 174 | s1 | 174 |
| V5 | s4 | 1,309 | s7 | 12,360 | WT | 5,198 | s1 | 174 |
| V6 | s4 | 1,309 | WT | 4,947 | WT | 5,198 | WT | 5,198 |
| V7 | WT | 5,896 | WT | 4,947 | WT | 5,198 | s9 | 78,824 |

**Table S6 Top 10 enriched variants in the 4-site fraction of the RBS oriPL library after round 4.**

| ***plsB* RBS sequence** | ***plsC* RBS sequence** | ***cdsA* RBS sequence** | ***pssA* RBS sequence** | ***plsB* RBS TIR** | ***plsC* RBS TIR** | ***cdsA* RBS TIR** | ***pssA* RBS TIR** | **Number of reads** |
|---|---|---|---|---|---|---|---|---|
| TTTAAGGAGGT AAAATCCAT | TTTAAGGAGAA TATATACAT | GATAAGGAGGT GATATACAT | TTTAAGAAGGA GGTATACAT | 72,247 | 12,360 | 78,824 | 31,879 | 181 |
| TTTAAGGAGGT AAAATCCAT | TTTAAGGCGAA TATATACAT | GATAAGGAGGT GATATACAT | TTTAAGAAGGA GGTATACAT | 72,247 | 249 | 78,824 | 31,879 | 172 |
| TTTAAGGAGGT AAAATCCAT | TTTAAGGGGAA TAGATATAT | TTTAAGAAGGA GGTATACAT | GATAAGGAGGT GATATACAT | 72,247 | 938 | 31,879 | 78,824 | 162 |
| TTTAAGGAGGT AAAACCCAT | TTTAAGGGGAA TAGATATAT | TTTAAGAAGGA GGTATACAT | GATAAGGAGGT GATATACAT | 21,409 | 938 | 31,879 | 78,824 | 18 |
| TTTAAGGAGGT AAAATCCAT | TTTAAGGGGAA TAGATACAT | TTTGAGAAGGA GATATACAT | GATAAGGAGGT GATATACAT | 72,247 | 3,327 | 4,794 | 78,824 | 17 |
| TTTAAGGAGGT AAAATCCAT | TTTAAGGGGAA TATATACAT | TTTAAGAAGGA GGTATACAT | GATAAGGAGGT GATATACAT | 72,247 | 5,725 | 31,879 | 78,824 | 16 |
| TTTAAGGAGGT AAAACCCAT | TTTAAGGCGAA TATATACAT | GATAAGGAGGT GATATACAT | TTTAAGAAGGA GGTATACAT | 21,409 | 249 | 78,824 | 31,879 | 15 |
| TTTAAGGAGGT AAAACCCAT | TTTAAGGGGAA TATATACAT | TTTAAGAAGGA GGTATACAT | GATAAGGAGGT GATATACAT | 21,409 | 5,725 | 31,879 | 78,824 | 14 |
| TTTAAGGAGGT AAAACCCAT | TTTAAGGGGAA TAGATACAT | TTTGAGAAGGA GATATACAT | GATAAGGAGGT GATATACAT | 21,409 | 3,327 | 4,794 | 78,824 | 13 |
| TTTAAGGAGGT AAAACCCAT | TTTAAGGAGAA TATATACAT | TTTAAGAAGGA GGTATACAT | GATAAGGAGGT GATATACAT | 21,409 | 12,360 | 31,879 | 78,824 | 11 |

**Table S7 Calculation of FACS screening coverage for the theoretical GC library.** Calculation based on 4,500,000 screened events and 13% PS-producing liposomes (for wild type). With a MOSAIC editing efficiency of ~50% per target locus, we obtained a library that consisted for 9% of variants with all target loci mutated, and the remainder of the library consisted of library fractions where zero, one, two or three loci were mutated (**Fig. S11a**). Using nanopore sequencing, we detected 565, 9,236, 7,785 and 2,922 library variants with one, two, three and four mutated RBSs, respectively, in the initial library. This corresponded to 99%, 8%, 0.1% and 0.001% of all theoretical library variants with one, two, three and four mutated RBSs, respectively. Potentially, more of the theoretical library is covered but not detected with the current sequencing depth.

| **Screening coverage for DNA variants:** | | | | |
|---|---|---|---|---|
| **Library fraction** | **Number of mutated loci** | **Number of DNA variants** | **Fraction of total library** | **Per-variant screening coverage** |
| 1 | 0 (WT) | 1 | 0.14 | 81,900 |
| 2 | 1 | 572 | 0.25 | 256 |
| 3 | 2 | 121,158 | 0.29 | 1 |
| 4 | 3 | 11,290,300 | 0.23 | 0.01 |
| 5 | 4 | 391,241,153 | 0.09 | 0.0001 |
| **Screening coverage for %GC variants:** | | | | |
| **Library fraction** | **Number of mutated loci** | **Number of %GC variants** | **Fraction of total library** | **Per-variant screening coverage** |
| 1 | 0 (WT) | 1 | 0.14 | 81,900 |
| 2 | 1 | 6 | 0.25 | 24,375 |
| 3 | 2 | 36 | 0.29 | 4,713 |
| 4 | 3 | 216 | 0.23 | 623 |
| 5 | 4 | 1,296 | 0.09 | 41 |

**Table S8 Most enriched variants (variant frequency > 0.02) of the GC oriPL library after round 4.**

| **Gene** | **DNA sequence** | **Predicted TIR (a.u.)** |
|---|---|---|
| *plsB* | CTTTTGTTATCCA | 5,870 |
| *plsB* | ATTTTGCTATCCA | 20,695 |
| *plsB* | CTTTTGTTACCCC | 5,870 |
| *plsC* | ATATATCTTTCGC | 8,605 |
| *plsC* | TTATATATTCCGG | 34,414 |
| *plsC* | CTACATTTTCCGA | 21,844 |
| *plsC* | ATATATCTTCCGA | 2,975 |
| *plsC* | ATATATCTTTCGA | 4,859 |
| *plsC* | ATATATCTTTCGG | 6,809 |
| *cdsA* | CAGCATTACCGT | 13,195 |
| *cdsA* | CAGCATGACCGT | 10,073 |
| *cdsA* | TAGTATTACTGG | 13,618 |
| *cdsA* | CAGCATTACCGT | 13,195 |
| *cdsA* | CAGCATGACCGT | 10,073 |
| *pssA* | TAGTATTACGGT | 5,611 |
| *pssA* | TAGCATTACAGG | 14,700 |

**Table S9 Clonal DNA variants from the oriPL GC library.**

| | ***plsB* RBS** | | | ***plsC* RBS** | | |
|---|---|---|---|---|---|---|
| | **Sequence** | **%GC** | **Predicted TIR** | **Sequence** | **%GC** | **Predicted TIR** |
| WT | WT (TTTCTGCTATCCT) | 0.38 | 5,896 | WT (ATATATCTTTCGT) | 0.23 | 4,947 |
| V8 | ATTTTGCTATCCA | 0.31 | 20,695 | ATATATCTTTCGC | 0.31 | 8,605 |
| V9 | ATTTTGCTATCCA | 0.31 | 20,695 | ATATATCTTTCGC | 0.31 | 8,605 |
| V10 | ATTTTGCTATCCA | 0.31 | 20,695 | ATATATCTTTCGC | 0.31 | 8,605 |
| V11$ | ATTTTGCTATCCA | 0.31 | 12,389$ | ATATATCTTTCGC | 0.31 | 8,605 |

| | ***cdsA* RBS** | | | ***pssA* RBS** | | |
|---|---|---|---|---|---|---|
| | **Sequence** | **%GC** | **Predicted TIR** | **Sequence** | **%GC** | **Predicted TIR** |
| WT | WT (TAGCATGACTGG) | 0.50 | 5,198 | WT (TAGCATGACTGG) | 0.50 | 5,198 |
| V8 | WT | 0.50 | 5,198 | WT | 0.50 | 5,198 |
| V9 | CAGCATGACCGT | 0.58 | 10,073 | WT* | 0.50 | 9,289* |
| V10 | TAGTATTACTGT | 0.25 | 13,618 | TAGTATTACTGT | 0.25 | 13,618 |
| V11 | CAGCATGACCGT | 0.58 | 10,073 | CAGCATGACCGT | 0.58 | 10,073 |

$ V11 has a deletion at the 22nd nucleotide upstream the start codon of the *plsB* gene.

* V9 has 4 deletions 17-20 nucleotides upstream the start codon of the *pssA* gene.

**Table S10 List of plasmids.**

| Plasmid name | Description |
|---|---|
| G555 | Expression of PlsB, PlsC, CdsA and PssA under the control of T7 promoters. The 4-gene insert is flanked by Phi29 replication origins. Addgene #216483. |
| G435 | Expression of DNAP and TP under the control of T7 promoters. The 2-gene insert is flanked by Phi29 replication origins. |
| pORTMAGE-Ec1 | Enables high-efficiency plasmid and genomic recombineering in *E. coli*. Expresses single-stranded DNA annealing protein (SSAP) CspRecT and the dominant negative *E. coli* MutL mutant (EcMutL$^{E32K}$), controlled by XylS-Pm expression system. Addgene #138474. |

**Table S11 List of DNA oligos.** '*' indicates phosphorothioate bond. Red color indicates degenerate nucleotides.

| Primer name | Sequence (5' to 3') | Description |
|---|---|---|
| DNAP DEG | C*A*CAACGGTTTCCCTCTAGAAATAATTTTGTTTAACTTTAADAMGGAGSTATACATATGCCGCGTAAAATGTACAGCTGCGATTTTGAAAC | G435 RBS library<br>Degenerate MOSAIC oligo for mutating the *DNAP* RBS in plasmid G435. |
| TP DEG | G*A*CACCAGAGGGTTTACATGTTTATTTGTTTAACTTTAAGAAGSRGBTTGACTAATGGCACGCAGCCCGCGCATCCGCATCAAAGATAACG | G435 RBS library<br>Degenerate MOSAIC oligo for mutating the *TP* RBS in plasmid G435. |

***Table S11 continued***

| | | |
|---|---|---|
| TP_1 custom | G*A*CACCAGAGGGTTTACATGTT TATTTGTTTAACTTTAAGAAGGAG GTTTACTAATGGCACGCAGCCCG CGCATCCGCATCAAAGATAACG | G435 RBS library<br>Additional MOSAIC oligo for mutating the *TP* RBS in plasmid G435. This oligo was mixed with the degenerate *TP* oligo set before addition to the electroporation sample. |
| 1585 ChD | C*A*CAACGGTTTCCCTCTAGAAA TAATTTTGTTTAACTTTAAGAAGG AGGTATACATATGCCGCGTAAAA TGTACAGCTGCGATTTTGAAAC | G435 RBS variant (s10) targeting the *DNAP* RBS (MOSAIC oligo). |
| 1586 ChD | C*A*CAACGGTTTCCCTCTAGAAA TAATTTTGTTTAACTTTAATAAGG AGGTATACATATGCCGCGTAAAA TGTACAGCTGCGATTTTGAAAC | G435 RBS variant (s12) targeting the *DNAP* RBS (MOSAIC oligo). |
| 1587 ChD | C*A*CAACGGTTTCCCTCTAGAAA TAATTTTGTTTAACTTTAATACGG AGCTATACATATGCCGCGTAAAA TGTACAGCTGCGATTTTGAAAC | G435 RBS variant (s3) targeting the *DNAP* RBS (MOSAIC oligo). |
| 1588 ChD | C*A*CAACGGTTTCCCTCTAGAAA TAATTTTGTTTAACTTTAAGAAGG AGCTATACATATGCCGCGTAAAA TGTACAGCTGCGATTTTGAAAC | G435 RBS variant (s5) targeting the *DNAP* RBS (MOSAIC oligo). |
| 1589 ChD | C*A*CAACGGTTTCCCTCTAGAAA TAATTTTGTTTAACTTTAATAAGG AGATATACATATGCCGCGTAAAA TGTACAGCTGCGATTTTGAAAC | G435 RBS variant targeting the *DNAP* RBS (MOSAIC oligo). |
| 1590 ChD | G*A*CACCAGAGGGTTTACATGTT TATTTGTTTAACTTTAAGAAGGAG GTTGACTAATGGCACGCAGCCCG CGCATCCGCATCAAAGATAACG | G435 RBS variant (s12) targeting the *TP* RBS (MOSAIC oligo). |
| 1591 ChD | G*A*CACCAGAGGGTTTACATGTT TATTTGTTTAACTTTAAGAAGCGG GTTGACTAATGGCACGCAGCCCG CGCATCCGCATCAAAGATAACG | G435 RBS variant (s3) targeting the *TP* RBS. |
| 1592 ChD | G*A*CACCAGAGGGTTTACATGTT TATTTGTTTAACTTTAAGAAGCAG TTTGACTAATGGCACGCAGCCCG CGCATCCGCATCAAAGATAACG | G435 RBS variant (s4) targeting the *TP* RBS (MOSAIC oligo). |
| PlsB DEG | C*A*ACGGTTTCCCTCTAGAAATA ATTTTGTTTAACTTTAAGGAGGKA AAMBCCATATGACTTTCTGCTAT CCTTGCCGCGCATTTGCATTA | G555 RBS library<br>Degenerate MOSAIC oligo for mutating the *plsB* RBS in plasmid G555. |
| PlsC DEG | C*A*ACGGTTTCCCTCTAGAAATA ATTTTGTTTAACTTTAAGGVGAAT ADATACATATGCTATATATCTTTC GTCTTATTATTACCGTGATT | G555 RBS library<br>Degenerate MOSAIC oligo for mutating the *plsC* RBS in plasmid G555. |
| PssA_CdsA DEG | A*T*AACAATTCCCCTCTAGAAAT AATTTTGTTTAACTTTAAGARKGA GKTATACATATGGCTAGCATGAC TGGTGGACAGCAAATGGGTCG | G555 RBS library<br>Degenerate MOSAIC oligo for mutating the *cdsA and pssA* RBSs in plasmid G555. |

***Table S11 continued***

| | | |
|---|---|---|
| PlsB custom | C*A*ACGGTTTCCCTCTAGAAATAATTTTGTTTAACTTTAAGGAGGTAATATACATATGACTTTCTGCTATCCTTGCCGCGCATTTGCATTA | G555 RBS library<br>Additional MOSAIC oligo for mutating the *plsB* RBS in plasmid G555. This oligo was mixed with the degenerate *plsB* oligo set before addition to the electroporation sample. |
| PlsC_1 custom | C*A*ACGGTTTCCCTCTAGAAATAATTTTGTTTAACTTTAAGGAGGTAATATACATATGCTATATATCTTTCGTCTTATTATTACCGTGATT | G555 RBS library<br>Additional MOSAIC oligo for mutating the *plsC* RBS in plasmid G555. This oligo was mixed with the degenerate *plsC* oligo set before addition to the electroporation sample. |
| PlsC_2 custom | C*A*ACGGTTTCCCTCTAGAAATAATTTTGTTTAACTTTAAGGAGGTAAGATACATATGCTATATATCTTTCGTCTTATTATTACCGTGATT | G555 RBS library<br>Additional MOSAIC oligo for mutating the *plsC* RBS in plasmid G555. This oligo was mixed with the degenerate *plsC* oligo set before addition to the electroporation sample. |
| PssA_CdsA custom | A*G*CGGATAACAATTCCCCTCTAGAAATAATTTTGTTTAACGATAAGGAGGTGATATACATATGGCTAGCATGACTGGTGGACAGCAAATG | G555 RBS library<br>Additional MOSAIC oligo for mutating the *cdsA and pssA* RBSs in plasmid G555. This oligo was mixed with the degenerate *cdsA/pssA* oligo set before addition to the electroporation sample. |
| 1543 ChD | C*A*ACGGTTTCCCTCTAGAAATAATTTTGTTTAACTTTAAGGAGGTAAAATCCATATGACTTTCTGCTATCCTTGCCGCGCATTTGCATTA | G555 RBS variant (s12) targeting the *plsB* RBS (MOSAIC oligo). |
| 1544 ChD | C*A*ACGGTTTCCCTCTAGAAATAATTTTGTTTAACTTTAAGGAGGGAAAACCCATATGACTTTCTGCTATCCTTGCCGCGCATTTGCATTA | G555 RBS variant (s4) targeting the *plsB* RBS (MOSAIC oligo). |
| 1545 ChD | C*A*ACGGTTTCCCTCTAGAAATAATTTTGTTTAACTTTAAGGAGAATATATACATATGCTATATATCTTTCGTCTTATTATTACCGTGATT | G555 RBS variant (s7) targeting the *plsC* RBS (MOSAIC oligo). |
| 1547 ChD | C*A*ACGGTTTCCCTCTAGAAATAATTTTGTTTAACTTTAAGGGGAATAGATATATATGCTATATATCTTTCGTCTTATTATTACCGTGATT | G555 RBS variant targeting the *plsC* RBS (MOSAIC oligo). |
| 1549 ChD | A*T*AACAATTCCCCTCTAGAAATAATTTTGTTTAACTTTAAGAATGAGTTATACATATGGCTAGCATGACTGGTGGACAGCAAATGGGTCG | G555 RBS variant (s1) targeting the *cdsA/pssA* RBSs (MOSAIC oligo). |
| 1481 ChD | A*T*AATTTTGTTTAACTTTAAGAAGGAGATATACATATGACNTTYTGYTAYCCNTGCCGCGCATTTGCATTATTAACCAGAGGCTTTACAT | G555 GC library<br>Degenerate MOSAIC oligo for modification of the GC content in the first five codons after the start codon of the *plsB* gene in plasmid G555. |
| 1482 ChD | A*T*AATTTTGTTTAACTTTAAGAAGGAGATATACATATGCTNTAYATHTTYCGNCTTATTATTACCGTGATTTACAGCATCTTAGTCTGTG | G555 GC library<br>Degenerate MOSAIC oligo for modification of the GC content in the first five codons after the start codon of the *plsC* gene in plasmid G555. |

***Table S11 continued***

| | | |
|---|---|---|
| 1483 ChD | T*C*TAGAAATAATTTTGTTTAACTTTAAGAAGGAGATATACATATGGCNAGYATKACNGKTGGACAGCAAATGGGTCGCGGATCCGGCTGC | G555 GC library<br>Degenerate MOSAIC oligo for modification of the GC content in the first five codons after the start codon of the T7 tag preceding the *cdsA* and *pssA* genes in plasmid G555. |
| 1551 ChD | A*T*AATTTTGTTTAACTTTAAGAAGGAGATATACATATGACATTTTGCTATCCATGCCGCGCATTTGCATTATTAACCAGAGGCTTTACAT | G555 GC variant targeting *plsB* (MOSAIC oligo). |
| 1554 ChD | A*T*AATTTTGTTTAACTTTAAGAAGGAGATATACATATGCTATATATCTTTCGCCTTATTATTACCGTGATTTACAGCATCTTAGTCTGTG | G555 GC variant targeting *plsC* (MOSAIC oligo). |
| 1557 ChD | T*C*TAGAAATAATTTTGTTTAACTTTAAGAAGGAGATATACATATGGCCAGCATGACCGTTGGACAGCAAATGGGTCGCGGATCCGGCTGC | G555 GC variant targeting *cdsA/pssA* (MOSAIC oligo). |
| 1560 ChD | T*C*TAGAAATAATTTTGTTTAACTTTAAGAAGGAGATATACATATGGCTAGTATTACTGTTGGACAGCAAATGGGTCGCGGATCCGGCTGC | G555 GC variant targeting *cdsA/pssA* (MOSAIC oligo). |
| 491 ChD | 5'Phos-AAAGTAAGCCCCCACCCTCACATG | PCR primer binding to oriL191 in G435 and G555. |
| 492 ChD | 5'Phos-AAAGTAGGGTACAGCGACAACATACAC | PCR primer binding to oriR194 in G435 and G555. |
| 1523 ChD | 5'Phos-AAAGTAAGCCCCCACCCTCACATGATACC | Alternative PCR primer for 491 ChD. |
| 1524 ChD | 5'Phos-AAAGTAGGGTACAGCGACAACATACACCATTTCC | Alternative PCR primer for 492 ChD. |
| 1459 ChD | CTCCTAATATCGACATAATCCGTCGATCCTCG | PCR primer binding to oriL191 used for recovery of oriPL from liposomes after FACS. |
| 1460 ChD | CCCCATTGACCGACTATCTTCGACAAG | PCR primer binding to oriR194 used for recovery of oriPL from liposomes after FACS. |
| 1301 ChD | 5'Phos-AAAGTAAGCCCCCACCCTCACATGATACCATTCTCCTAATATCGACATAATCCGTCGATCC | PCR primer binding to oriL191 used for recovery of oriPL from liposomes after FACS. |
| 1518 ChD | 5'Phos-AAAGTAGGGTACAGCGACAACATACACCATTTCCCCATTGACCGACTATCTTCG | PCR primer binding to oriR194 used for recovery of oriPL from liposomes after FACS. |
| 976 ChD | GGATGAAGACTACCCGCTGC | qPCR primer binding in *DNAP*. |
| 977 ChD | ACAGGTCTGCGATTTCACCG | qPCR primer binding in *DNAP*. |

**Table S12** Total number of reads of the different library fractions (n-site fractions) for the RBS library of the phospholipid synthesis pathway, supplementing the data in **Fig. S12**:

| **Round** | **1-site** | **2-site** | **3-site** | **4-site** |
|---|---|---|---|---|
| **R0** | 11,977 | 15,586 | 13,835 | 6,666 |
| **R1** | 19,404 | 17,740 | 14,456 | 7,134 |
| **R2** | 11,897 | 13,071 | 5,189 | 2,030 |
| **R3** | 9,528 | 7,790 | 3,268 | 970 |
| **R4** | 9,123 | 8,510 | 3,119 | 697 |

**Table S13** Total number of reads for the different library fractions of the GC library of the phospholipid synthesis pathway, supplementing the data in **Fig. S16**.

| **Round** | **1-site** | **2-site** | **3-site** | **4-site** |
|---|---|---|---|---|
| **R0** | 10,314 | 12,067 | 9,410 | 3,607 |
| **R1** | 37,127 | 32,459 | 17,614 | 5,850 |
| **R2** | 55,764 | 31,071 | 14,882 | 2,988 |
| **R3** | 41,912 | 34,681 | 12,152 | 1,309 |
| **R4** | 34,025 | 39,306 | 17,120 | 1,639 |
